# Augment rice *ANNEXIN* expression to counter planthopper *Nl*Annexin-like5 as an anti-virulence strategy against a major crop pest

**DOI:** 10.1101/2025.08.18.670976

**Authors:** Xiao-Ya Zhang, Shaoqin Li, Comzit Opachaloemphan, Chuan-Xi Zhang, Sheng Yang He, Yanjuan Jiang

## Abstract

The brown planthopper (BPH) is the most devastating insect pest in rice, posing a serious threat to global rice production. One attractive control strategy would be based on the understanding of the virulence mechanisms of BPH at the molecular level and then designing targeted methods to neutralize such mechanisms. Salivary proteins of BPH are important players in mediating rice-BPH interactions. Here, we describe a pivotal role of a watery saliva protein, *Nl*Annexin-like5 (*Nl*ANX5), in the rice-BPH interaction. RNA interference (RNAi) of *NlANX5* greatly compromised BPH feeding performance and survival rate on rice plants. *NlANX5*-RNAi BPH triggered a rapid calcium ion influx in rice cells. The feeding and survival defects of *NlANX5*-RNAi BPH can be restored in *NlANX5*-expressing transgenic rice plants. *Nl*ANX5 targets rice annexin (*Os*ANN) proteins, including *Os*ANN2 and *Os*ANN8. Further analysis with *Nl*ANX5 and *Os*ANN2 as well as *Os*ANN8 showed that *Nl*ANX5 displaces *Os*ANN2 and *Os*ANN8 from the rice cell membrane. The *osann2 osann8* mutant rice plants are hypersusceptible to BPH infestation. In contrast, enhanced expression of *OsANN2* and *OsANN8* gene resulted in robust rice resistance against BPH. This study highlights a successful example of identifying and augmenting the expression of the host targets of a major BPH virulence effector as a promising anti-virulence strategy against an important crop pest.

**Significance Statement:** Insect pests pose a serious biotic threat to crop production worldwide. Understanding how insect pests attack plants could inspire innovative pest control measures to enhance global food security. The brown planthopper (BPH; *Nilaparvata lugens* Stål) is the most devastative insect pest in rice. In this study, we discovered that BPH secretes a salivary protein, *Nl*Annexin-like5 (*Nl*ANX5), to target rice host annexins that are associated with calcium fluxes and activation of multiple rice defense pathways. This knowledge led to employing enhanced expression of rice annexin-encoding genes to successfully defeat the virulence function of *Nl*ANX5. Results have significant implications in the development of anti-virulence breeding strategies against BPH.

## Introduction

The molecular wars between plants and herbivores have led to evolution of sophisticated plant defensive mechanisms as well as herbivore countermeasures, which provide a rich context to understand how host-herbivore factors collectively shape their interactions. Among herbivores of major agricultural importance are various piercing-sucking insects, of which *Nilaparvata lugens* Stål (Brown planthopper, BPH) is one of the most harmful pests (1−5). BPH can cause billions of tons of global rice loss yearly and, in recent years, has emerged as a model for understanding plant-insect interactions. To date, more than 30 rice BPH resistance genes have been identified in the rice genome, some of which, along with chemical pesticides, are used as major weapons to control BPH (2). Unfortunately, BPH evolves new biotypes rapidly that defeat many *R* genes. Therefore, development of novel control strategies for BPH is urgently needed. One alternative strategy would be based on the understanding of the virulence mechanisms of BPH at the molecular level and then designing targeted methods to neutralize such mechanisms.

During feeding, sap-sucking insects activate a plethora of plant defense responses, including calcium (Ca^2+^) fluxes, reactive oxygen species (ROS) bursts, and defense hormone production and signaling (6−9). For example, many studies have shown a crucial role of Ca^2+^ in regulating plant immune response against pathogen (10−13). The role of Ca^2+^ in plant-insect interaction has also attracted the attention of researchers in recent years. Wounding caused by caterpillar feeding triggered a rapid [Ca^2+^]_cyt_ increase at the herbivory site to activate jasmonate (JA) defense responses in distal leaves (14). It was shown that the glutamate receptor-like (GLRs) proteins in *Arabidopsis thaliana* mediate Ca^2+^ response to wounding and insect attack, which activate JA-mediated defense responses (14). Ca^2+^ responses are also integral to other branches of the plant immune system, including salicylic acid defense pathway, pattern triggered immunity and effector-triggered immunity (15−16). Interestingly, calcium-binding salivary proteins have been identified in certain piercing-sucking insects. For instance, Ca^2+^-binding proteins secreted by the aphid *Megoura viciae* and the green rice leafhopper *Nephotettix cincticeps* prevented sieve element occlusion and facilitated continuous insect feeding from sieve tubes (17−18). A salivary *EF-hand* protein (*NlSEF1*) from BPH suppressed hydrogen peroxide (H_2_O_2_) production in rice (19). However, the underlying molecular mechanism by which these insect effectors modulate plant Ca^2+^ oscillation remains unclear, nor have approaches to inactivate these insect virulence effectors been reported.

Annexins are Ca^2+^-associated proteins and, in their Ca^2+^-bound form, bind to the membrane (20−21). Annexins are found in plants, animals and microorganisms (22). Plant annexins have been implicated in pathogen and herbivorous responses. For example, *Os*ANN1 has recently been shown to negatively regulate blast disease resistance in rice through inactivating JA signaling (23). An ANNEXIN-like protein of the cereal cyst nematode *Heterodera avenae* suppresses plant defense (24). However, the underlying mechanism remains unknown.

Here, we report that BPH delivers an annexin protein, *Nilaparvata lugens* Annexin-like5 (*Nl*ANX5), into rice tissues to suppress rapid increases of cytosolic Ca^2+^ concentration. Mechanistically, *Nl*ANX5 form heterodimers with rice host annexins, particularly *Os*ANN2 and *Os*ANN8, which positively regulate multiple rice defenses against BPH. The identification of rice annexins as the host targets of *Nl*ANX5 led to transgenic expression of *OsANN2* and *OsANN8* as an effective control strategy to counter BPH infestation. Thus, this study not only revealed the molecular action of one of the most important virulence effectors of BPH, but also led to a promising anti-virulence breeding strategy to control a major crop pest.

## Results

### *Nl*ANX5 is secreted into rice sheath tissues during BPH feeding

In a previous study, we found that *NlANX5* was present in the watery saliva of BPH fed on artificial diet (25), suggesting that *NI*ANX5 might be an effector protein delivered into rice tissues during BPH feeding. To test this possibility, we characterized *NlANX5* further in this study. First, we examined the expression level of *NlANX5* in BPH salivary glands by RNA *in situ* hybridization, which showed the *NlANX5* was mainly expressed in the principal glands (PGs) and accessory glands (AGs) (Fig. 1*A*). The PGs of BPH are bilaterally paired structures that are primarily responsible for synthesizing and secreting effector proteins critical for insect-host interactions. AGs are smaller, paired structures positioned near the principal glands and contribute to saliva composition by secreting mucopolysaccharides and lubricants that facilitate stylet penetration into plant tissues (26, 27). The glandular epithelium contains specialized “A-follicles”, tubular secretory units that store and release saliva components during feeding (25).

**Fig. 1.**
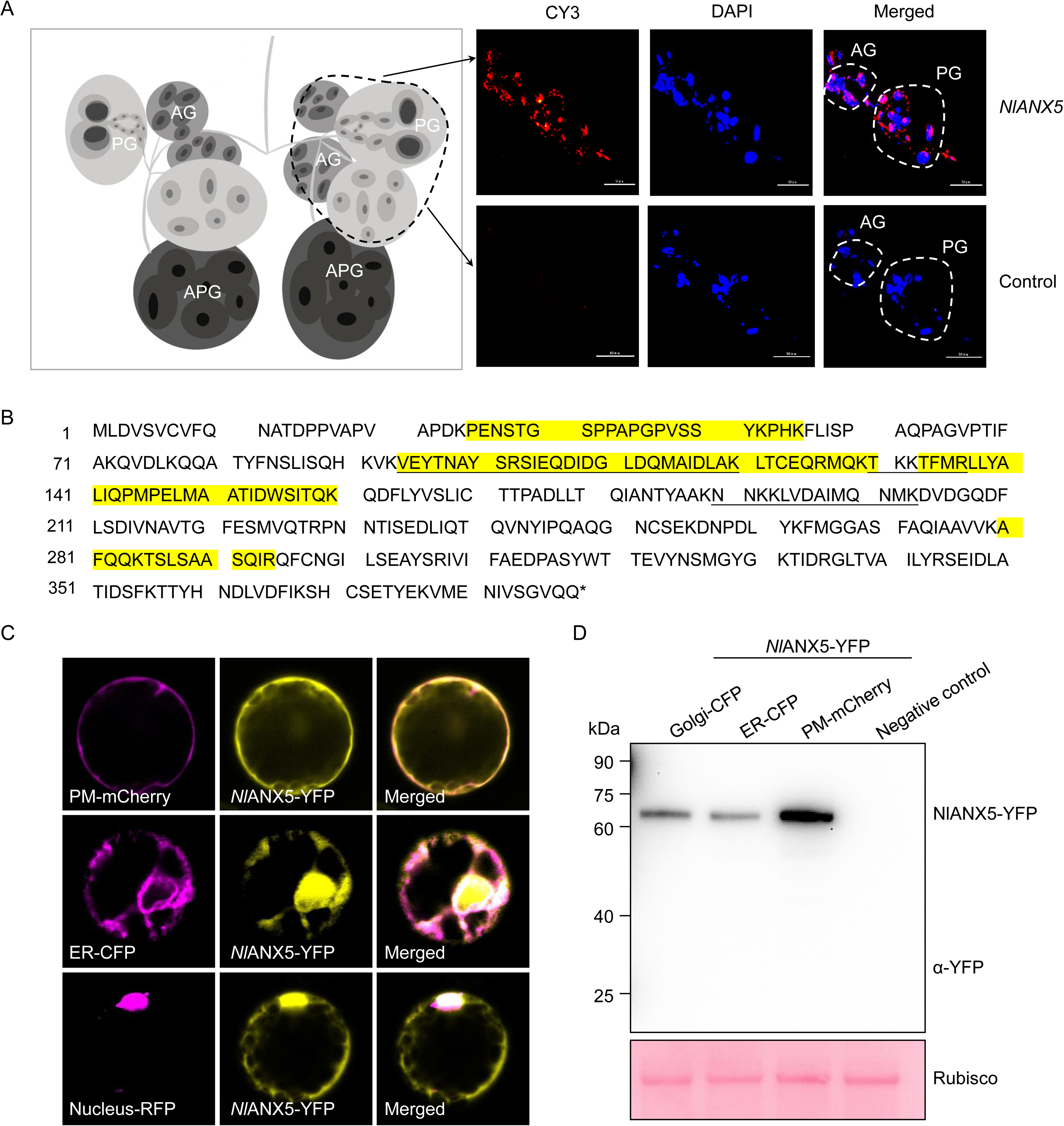
Characterization of *Nilaparvata lugens* Annexin-like5 (*Nl*ANX5). (*A*) RNA *in situ* hybridization of *NlANN5* in salivary glands isolated from 5^th^ instar BPHs (upper row). Sense probe was used as a negative control (lower row). PG, principal gland. AG, accessory gland. APG, a-follicle of the principal gland. (*B*) The amino acid sequence of *Nl*ANX5. The yellow highlighted amino acid residues indicate the peptides detected in BPH-fed rice sheath tissue by LC-MS analyses. The asterisk indicates the stop codon. The underlined amino acid residues indicate the peptides detected in the phloem exudate of BPH-infested rice by LC-MS analysis. (*C*) *Nl*ANX5 is localized in the plasma membrane (PM), endoreticulum (ER) and nucleus of rice cells. PM protein-mCherry and *Nl*ANX5-YFP fusion, ER protein-CFP and *Nl*ANX5-YFP or Nucleus-RFP and *Nl*ANX5-YFP proteins were co-expressed in rice protoplasts. The PM marker was the full-length Arabidopsis aquaporin 2A (AtPIP2A) (66), the ER marker was the signal peptide of Arabidopsis wall-associated kinase 2 (AtWAK2) (66), and the nucleus marker was the full-length Arabidopsis ELONGATED HYPOCOTYL 5 (AtHY5). See additional images in Fig. S2. (*D*) *Nl*ANX5-YFP fusion protein levels in rice protoplasts are detected with anti-GFP (Abmart). Protein samples were extracted from rice protoplasts co-expressing Golgi-CFP and *Nl*ANX5-YFP, ER-CFP and *Nl*ANX5-YFP, or PM-mCherry and *Nl*ANX5-YFP, respectively. Proteins from mock rice protoplasts were included as negative control. Ponceau S staining of the gel shows overall protein loading. Experiments were repeated three times with similar trends.

We next investigated if the *Nl*ANX5 protein is delivered to the rice tissue during BPH feeding by comparing rice leaf sheath protein profiles of cv. Nipponbare plants fed by BPH with those without BPH feeding using liquid chromatograph-mass spectrometer (LC-MS). We found four *Nl*ANX5-unique peptides in only BPH-fed leaf sheath tissue (Fig. 1*B*; *SI Appendix*, Fig. S1), consistent with the hypothesis that *Nl*ANX5 is secreted into the rice tissues by BPH during feeding. We further collected the phloem exudate from Nipponbare rice leaf sheaths, with or without BPH feeding, for LC-MS analysis. Three *Nl*ANX5-specific peptides were detected in the phloem exudate of Nipponbare plants that had been fed on by BPH (Fig. 1*B*), providing an independent line of evidence that *Nl*ANX5 is delivered into the phloem by BPH.

We next investigated the subcellular localization of *NI*ANX5 in rice leaf protoplasts transiently expressing *Nl*ANX5:YFP. We found that *Nl*ANX5:YFP showed a nonuniform distribution of YFP signal in the cell. Further colocalization studies with subcellular protein markers revealed that *NI*ANX5 was colocalized in the cytoplasm, various membranes and the nucleus (Fig. 1*C* and 1*D*; *SI Appendix*, Fig. S2).

### Transgenic expression of *NlANX5* in rice restores feeding and survival of *NIANX5*-knockdown BPH on rice plants

To further test the hypothesis that *NI*ANX5 acts inside host tissues, we constructed *NlANX5*-expressing transgenic rice (cv. Nipponbare) plants (see Methods). Twenty-two independent transgenic lines were produced, and eight lines were found to robustly express the *NlANX5* transcript (Fig. 2*A*). *NIANX5*-expressing plants did not display noticeable growth alterations compared to Nipponbare plants (*SI Appendix*, Fig. S3) and they were propagated to the T3 generation. Two independent stable lines were used for further assays.

**Fig. 2.**
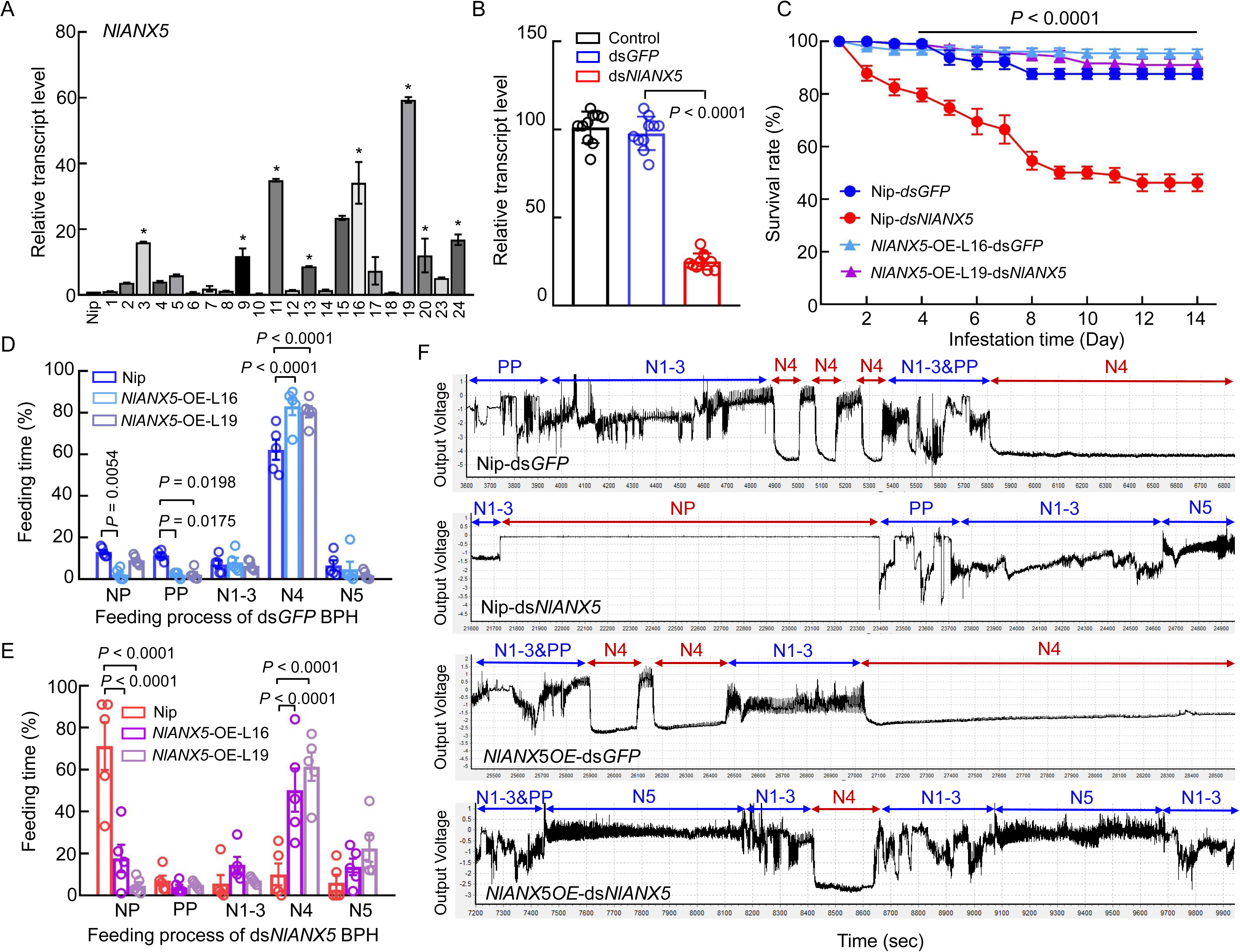
*Nl*ANX5 is critical for BPH’s survival on Nipponbare rice plants. (*A*) *NIANX5* transcript levels in the *NlANX5*-expressing transgenic rice plants. Nipponbare was used as a negative control for *NlANX5*-expressing plants, whereas the *OsACTIN* (*Os03g50885*) gene was used as an internal control for transcript quantification. Values are displayed as mean ± s.e.m. (two-way ANOVA, *n* = 3 biological replicates). Plant lines labeled with asterisks were found to robustly express the *NlANX5* transcript. (*B*) The relative gene expression of *NlANX5* at 3 dpi in *NlANX5*-RNAi and *GFP*-RNAi BPH. Wild type BPH was used as a control. Values are displayed as mean ± s.e.m. (two-way ANOVA, *n* = 30 biological replicates). (*C*) The survival rates of ds*GFP*- and ds*NlANX5*-injected BPH insects in Nipponbare (Nip) and *NlANX5-*OE rice plants. Values are displayed as mean ± s.e.m. (*n* = 8 biological replicates for each genotype). The black line on top indicates significant differences in survival rate between Nip and *NlANX5*-OE plants fed by ds*GFP* and ds*NlANX5* BPH (two-way ANOVA, *P* < 0.0001). The 2^nd^ instar nymphs were used in the survival assays. (*D* and *E*) Electrical penetration graph (EPG) monitoring of ds*GFP*- and ds*NlANX5*-treated BPH on Nip and *NlANX5*-OE rice plants. NP, nonpenetration. PP, pathway phase. N1–3, including N1 penetration initiation, N2 salivation and stylet movement and N3 extracellular activity near the phloem. N4, phloem sap ingestion. N5, xylem sap ingestion. Values are displayed as mean ± s.d. (two-way ANOVA, *n* ≥ 6 biological replicates). The adult BPH insects were used in the EPG assays. (*F*) The EPG waves recorded within 20 h feeding of ds*GFP* and ds*NlANX5* BPH on Nip and *NlANX5*-OE plants, as described in (*D* and *E*). Experiments were repeated three times with similar trends.

For BPH survival assay, double-stranded RNA of *NlANX5* (ds*NlANX5*) and the control green fluorescent protein gene (ds*GFP*) were injected into 1^st^ instar BPH nymphs following a previous study (28). Wild type BPH without RNAi treatment was used as an additional control. Quantitative real-time PCR (qPCR) analysis confirmed that the *NlANX5* transcript level was reduced to ∼25% in ds*NlANX5*-injected BPH compared to the wild type BPH control or ds*GFP*-injected BPH (Fig. 2*B*). ds*NlANX5* BPH performed poorly on wild-type Nipponbare plants, showing a lower survival rate compared with control ds*GFP* BPH, starting the second day post-feeding (Fig. 2*C*; *SI Appendix*, Fig. S4). Conversely, the Nipponbare rice seedlings fed by ds*NlANX5* BPH had fewer dead seedlings than those fed by ds*GFP* BPH (*SI Appendix*, Fig. S5*A*). The requirement of *NlANX5* for BPH survival in cv. Nipponbare is in line with a previous study conducted in the *japonica* cultivar Xiushui134 (25), suggesting that the function of *NlANX5* in promoting BPH survival is robust across different rice varieties. Importantly, *NlANX5*-expressing transgenic plants restored the survival of ds*NlANX5* BPH insects (Fig. 2*C*; *SI Appendix*, Fig. S4), providing further evidence that *NI*ANX5 functions inside the plant tissue to promote BPH survival in rice.

Next, we conducted BPH feeding behavior assay using the electrical penetration graph (EPG) method. EPG records the position, movement rates and duration of insect mouthparts within and near the host tissues (29−32). The BPH feeding process consists of five classic waveforms which include the non-penetration (NP) phase and the intracellular phase of activity in the phloem and the phloem sap ingestion phase (N4) (33−35). Compared to ds*GFP* BPH, ds*NlANX5* BPH spent a longer time in the NP phase and a shorter N4 feeding phase on wild type Nipponbare plants. Interestingly, *NlANX5*-expressing rice plants could complement the behavior defects of ds*NlANX5* BPH, including shortening the NP phase and greatly extending the N4 feeding phase (Fig. 2*D*−*F*).

We also conducted experiments to measure honeydew production during BPH feeding in wild-type Nipponbare and *NlANX5*-expressing rice plants. On Nipponbare plants ds*NlANX5* BPH had lower excrement of honeydew than ds*GFP* BPH, whereas ds*NlANX5* BPH fed on *NlANX5*-OE plants had almost the same amount of excrement as ds*GFP* BPH fed on wild-type Nipponbare plants (*SI Appendix* Fig. S5*E*). These results are consistent with a critical role of *Nl*ANX5 in sustaining normal BPH feeding behavior in rice plants.

### *Nl*ANX5 modulates cytosolic Ca^2+^ accumulation during BPH feeding

Annexins are phospholipid- and Ca^2+^-binding proteins (20). We wondered if *NI*ANX5 affects cytosolic Ca^2+^ homeostasis in rice cells during BPH feeding. To test this possibility, we generated transgenic rice (cv. Nipponbare) plants expressing a ratiometric calcium indicator, RGECO1-mTurquoise (hereinafter cyto-Ca^2+^-sensor) (*SI Appendix*, Fig. S6). We placed 20 5^th^ instar BPH on each cyto-Ca^2+^-sensor plant and captured the fluorescent signals within 24 min of BPH feeding. As shown in Fig. 3*A* and 3*B*, BPH feeding elevated the calcium level immediately. Notably, ds*NlANX5*-injected BPH triggered much higher fluorescent signals than ds*GFP*-injected BPH, suggesting *Nl*ANX5 plays an important role in dampening rapid cytosolic calcium increase during BPH feeding.

**Fig. 3.**
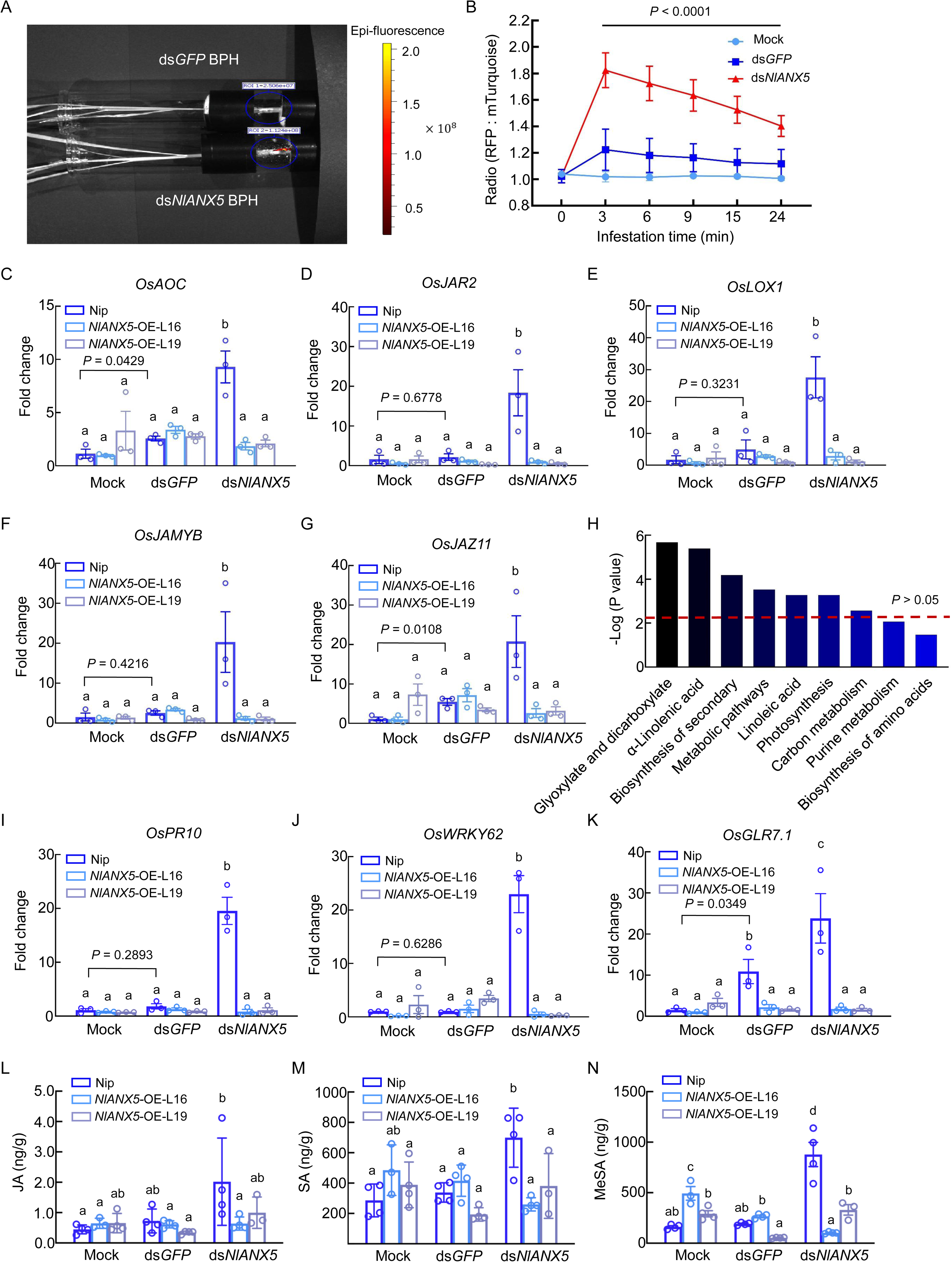
*Nl*ANX5 suppresses rapid Ca^2+^ response and expression of JA pathway genes. (*A*) Image at BPH feeding sites in Nipponbare plants expressing a ratiometric calcium indicator RGECO1-mTurquoise (cyto-Ca^2+^-sensor) captured by the In Vivo Imaging System (IVIS). Each sensor plant fed by 20 ds*NlANX5-*injected 5^th^ BPH nymphs or 20 ds*GFP*-injected BPH nymphs. (*B*) RFP:mTurquoise signal ratios within 24 min of ds*GFP* BPH or ds*NlANX5* BPH treatments compared to no BPH control. RFP was imaged at λEx = 532 nm and λEm = 580–640 nm, whereas mTurquoise was imaged at λEx = 405 nm and λEm = 460–520 nm. Values are displayed as mean ± s.e.m. (*n* = 3 biological replicates). The black line on top indicates significant differences in RFP: mTurquoise signal ratios between cyto-Ca^2+^-sensor plants fed by ds*GFP* and ds*NlANX5* BPH (two-way ANOVA, *P* < 0.0001). (*C–G*) Transcript levels of JA biosynthesis genes *OsAOC, OsLOX1* and *OsJAR2* and JA responsive genes *OsJAmyb* and *OsJAZ9* in Nip, *NlANX5*-OE-L16 and *NlANX5-OE-*L19 fed by ds*GFP* BPH or ds*NlANX5* BPH for 24 h. The *OsACTIN* (*Os03g50885*) gene was used as an internal control. (*H*) Main enrichment KEGG pathways of LC-MS analysis of leaf sheaths of Nip after BPH fed for 48 h. The significance of the enriched KEGG pathway is shown by *P* value calculated based on Fisher’s Exact Test. (*I–K*) Transcript levels of SA pathway genes, *OsPR10*, *OsWRKY62* and *OsGLR7.1*, in Nip and *NlANX5-OE* plants fed by 20 individual 5^th^ instar BPH for 24 h. The *OsACTIN* (*Os03g50885*) gene was used as an internal control. (*L–N*) Contents of defense hormones, JA, SA and MeSA, in Nip and *NlANX5-OE* plants. 3-week-old rice plants were fed by 20 individual 5^th^ instar BPH for 48 h. Values are displayed as mean ± s.e.m. (*n* = 3 biological replicates) in (*C–G*) and (*I–K*) and as mean ± s.d. (*n* = 4 biological replicates) in (*L–N*). Different letters indicate statistically significant differences analyzed by two-way ANOVA (Tukey test, *P* < 0.05). Mock group in (*C–G*) and (*I–N*) indicates plants infested by BPH without injection. Experiments were repeated three times.

### *Nl*ANX5 prevents activation of multiple defense pathway

A rapid influx of Ca^2+^ into the cytosol (Ca^2+^ spikes) is one of the major early plant immune responses (36−38). Recent studies showed that wound-induced Ca^2+^ spikes lead to JA gene expression in Arabidopsis (14, 39−41). Because JA signaling is associated with plant immune response to insect herbivore attacks (42), we examined the possibility that *Nl*ANX5 suppression of the Ca^2+^ burst during BPH feeding may modulate JA signaling pathway of rice plants. We first conducted KEGG analysis of differentially expressed proteins identified in LC-MS analysis of leaf sheath fed by BPH (see Methods), which revealed that proteins involved in the biosynthesis of α-linolenic acid, the precursor of JA-Ile biosynthesis, is significant enrichment after BPH fed for 48 hours in Nipponbare plants (Fig. 3*H*). Next, we examined the expression of JA biosynthesis genes *OsAOC*, *OsLOX1* and *OsJAR2* as well as JA signaling genes *OsJAMYB*, and *OsJAZ11* in Nipponbare and *NlANX5*-overexpressing plants. We found that JA pathway genes were up-regulated in Nipponbare plants during feeding by ds*NlANX5*-injected BPH (Fig. 3*C–G*). Notably, accumulation of JA as well as JA pathway gene expression were almost completely prevented during feeding by ds*NlANX5*-injected BPH in *NlANX5*-expressing plants (Fig. 3*C–G*, 3*L*).

However, the effect of *Nl*ANX5 is not restricted to JA pathway gene expression. Ca^2+^ response is also linked to SA defense pathway for example (43−44). Indeed, we found that *Nl*ANX5 prevents activation of SA pathway gene and *OsGLR* gene expression, including *OsPR10*, *OsWRKY62* and *OsGLR7.1* (Fig. 3*I–K*), and accumulation of SA and MeSA (Fig. 3*M* and 3*N*). This is consistent with results from BPH survival assay on Nipponbare and the *coi1a* mutant rice plants, which contains a loss-of-function JA receptor COI1. Although the *coi1a* mutant plants can partially restored the survival rate of ds*NlANX5* BPH (*SI Appendix*, Fig. S7), the restoration was not complete, suggesting that multiple rice pathways (i.e, in addition to JA pathway) are impacted by *Nl*ANX5.

### *Nl*ANX5 interacts with *Os*ANN proteins and displaces *Os*ANN2 and *Os*ANN8 from the host membrane

Next, we investigated how an insect-derived annexin could suppress the Ca^2+^ spike and downstream defense pathway activation during BPH feeding. Because rice has its own endogenous annexin proteins, one possibility, we reasoned, is that *NI*ANX5 directly targets and interferes with the function of rice annexin protein(s). To test this possibility, we performed co-immunoprecipitation assay between *NI*ANX5 and ten rice annexin proteins in *Nicotiana benthamiana* leaves. As shown in Fig. 4*A* and 4*B*, *Os*ANN2, 3 and 8 strongly co-immunoprecipitated with *Nl*ANX5-YFP, whereas *Os*ANN4, 5, 6 and 7 showed moderate interactions with *Nl*ANX5-YFP. No interaction between *NI*ANX5-YFP and *Os*ANN1 was detected. *Os*ANN9 and *Os*ANN10 were not reliably detectable in our experiments, which provents accurate assessment of their interactions with *Nl*ANX5 (Fig. 4*B*). To further validate these interactions, *Os*ANN2, *Os*ANN6 and *Os*ANN8 were selected for bimolecular fluorescence complementation (BiFC) assay (Fig. 4*C*), with *Os*ANN1 used as a non-interaction control. The BiFC results further confirm the physical interactions between *Nl*ANX5 and *Os*ANN2, *Os*ANN6 and *Os*ANN8 *in planta*. We additionally performed co-IP assays with or without Ca^2+^. As shown from Fig. 4*G* and 4*H*, Ca^2+^ could slightly enhance the recovery of *Os*ANN2 and *Os*ANN8 proteins in the co-IP experiments. This seems to be consistent with the prediction by AlphaFold3 (45), which shows higher confidence scores for *Nl*ANX5 interaction with *Os*ANN2, *Os*ANN6 and *Os*ANN8 in the presence of calcium ions (*SI Appendix* Table S1). Taken together, these results suggest that *Nl*ANX5 targets a subset of rice annexins, including at least *Os*ANN2, *Os*ANN6 and *Os*ANN8, in rice. We also examined subcellular locations of *NI*ANX5 and *Os*ANN proteins in rice protoplasts. *Nl*ANX5, *Os*ANN2 and *Os*ANN8 were each found to be co-localized with endoplasmic reticulum (ER)-, Golgi- and plasma membrane-associated protein markers (Fig. 1*C*; Fig. 4*D*; *SI Appendix*, Fig. S8), illustrating their broad subcellular distributions.

**Fig. 4.**
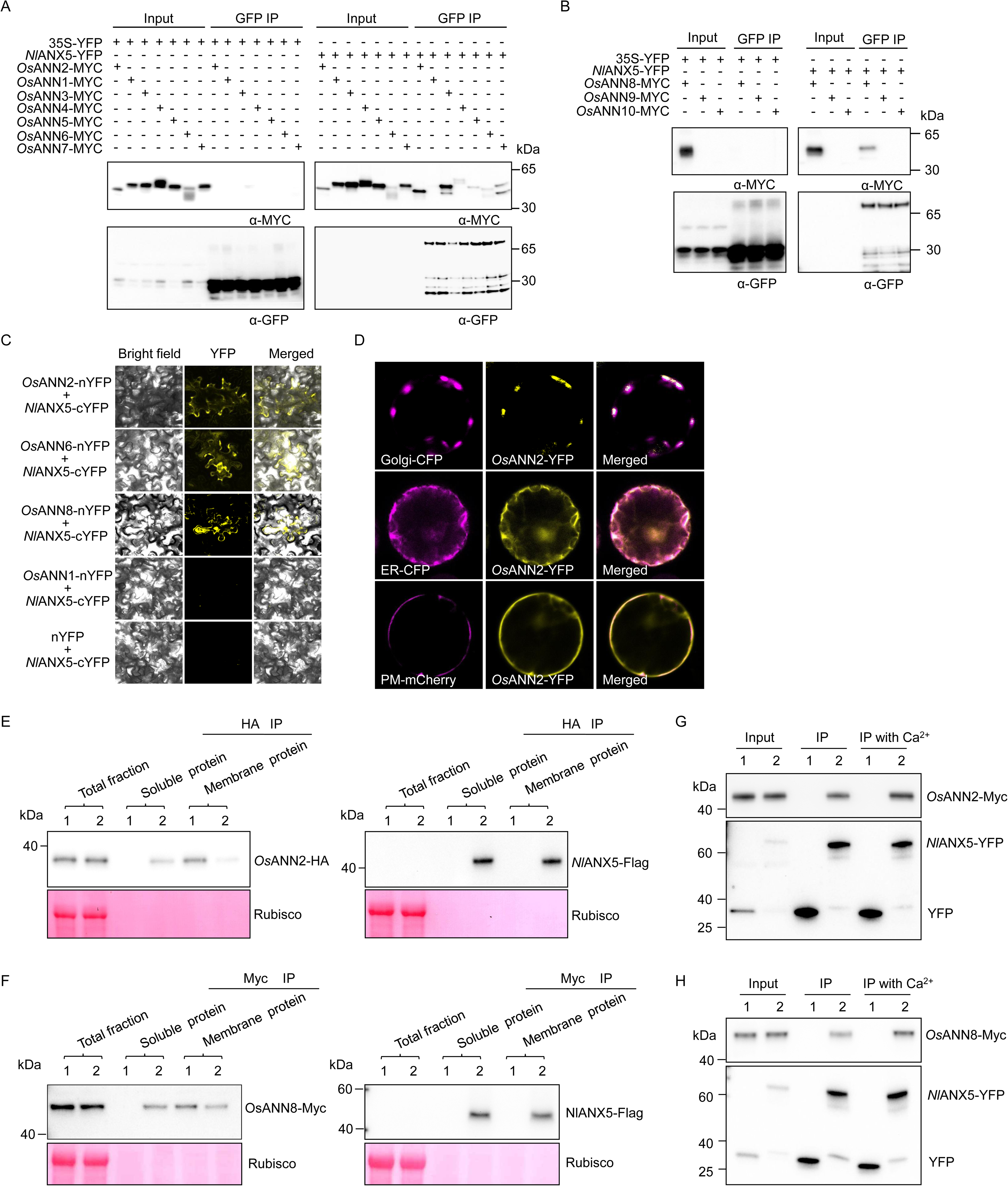
*Nl*ANX5 interacts with *Os*ANNs *in planta*. (*A–B*) *Nl*ANX5:YFP was co-expressed with individual *Os*ANN-MYC fusion proteins. Protein input for *Nl*ANX5:YFP and *Os*ANN-MYC was detected by anti-GFP antibody (no. M20004, Abmart) and anti-Myc antibody (no. M20002, Abmart), respectively. *Nl*ANX5-YFP was immunoprecipitated using the GFP-trap agarose (no. gta-20, ChromoTek). *Os*ANN proteins co-immunoprecipitated with *Nl*ANX5-GFP were detected by anti-Myc antibody. IP, immunoprecipitation; 35S:YFP samples were used as negative controls. (*C*) BiFC assay. Fluorescence was observed in the transformed *N. benthamiana* cells co-expressing *Nl*ANX5-cYFP with *Os*ANN2-nYFP, *Os*ANN6-nYFP, *Os*ANN8-nYFP, or *Os*ANN1-nYFP. *Os*ANN1 used as a non-interaction control, whereas *Nl*ANX5-YFP with nYFP used as negative control. (*D*) *Os*ANN2 is colocalized with the YFP signals in the Golgi (upper row), ER (middle row) and plasm membrane (PM) (lower row) of rice cells. Golgi-CFP and *Os*ANN2-YFP fusion protein, ER-CFP and *Os*ANN2-YFP, or PM-mCherry and *Os*ANN2-YFP fusion proteins were co-expressed in rice protoplasts 16 h after the corresponding DNA constructs were introduced into rice protoplasts via polyethylene glycol-mediated transformation. (*E–F*) *Nl*ANX5 displaced *Os*ANN2 and *Os*ANN8 from the host membrane. *Os*ANN2-HA, *Os*ANN8-Myc and *Nl*ANX5-Flag proteins were detected in different components of protein fractions by anti-HA, anti-Myc or anti-Flag antibodies. Sample 1 represents rice protoplasts expressed with *Os*ANN2-HA or *Os*ANN8-Myc, while sample 2 indicates rice protoplasts co-expressed with *Os*ANN2-HA and *Nl*ANX5-Flag or *Os*ANN8-Myc and *Nl*ANX5-Flag. Soluble and membrane proteins were immunoprecipitated by HA-trap, Myc-trap or Flag-trap beads, respectively, to detect *Os*ANN2-HA, *Os*ANN8-Myc and *Nl*ANX5-Flag proteins. (*G–H*) Ca^2+^ promotes interactions of *Nl*ANX5 with *Os*ANN2 and *Os*ANN8. *Nl*ANX5:YFP was co-immunoprecipitated with *Os*ANN2-Myc or *Os*ANN8-Myc fusion proteins without or with 5 mM Ca^2+^. Sample 1 represents rice protoplasts expressing YFP and *Os*ANN2-Myc or *Os*ANN8-Myc, whereas sample 2 contains rice protoplasts expressing *Nl*ANX5-YFP with *Os*ANN2-Myc or *Os*ANN8-Myc. *Nl*ANX5-YFP was immunoprecipitated using the GFP-trap agarose and detected by anti-GFP antibody. *Os*ANN2 or *Os*ANN8 proteins co-immunoprecipitated with *Nl*ANX5-YFP were detected by anti-Myc antibody. IP, immunoprecipitation. Experiments were repeated three times with similar trends.

Annexins interact with the membrane as part of their function (20, 46−47). To explore the mechanism of virulence action of *Nl*ANX5, we investigated a possible effect of *Nl*ANX5 on *Os*ANN interaction with rice membranes. For this purpose, we track membrane associations of *Os*ANN2, *Os*ANN6 and *Os*ANN8 in rice protoplasts with or without *Nl*ANX5. We found that, in the presence of *Nl*ANX5, *Os*ANN2 and *Os*ANN8 association with rice membranes was partially displaced and that an increasing amount of *Os*ANN2 and *Os*ANN8 was found in the cytosol (Fig. 4*E* and 4*F*). Interestingly, *Os*ANN6 association with rice membranes was not affect by *Nl*ANX5 (*SI Appendix*, Fig. S9), showing a certain degree of specificity in the membrane displacement of *Os*ANN2 and *Os*ANN8 by *Nl*ANX5.

### *Os*ANN2 and *Os*ANN8 positively regulate rice resistance against BPH

Because *Nl*ANX5 targets several *Os*ANN proteins (Fig. 4), we hypothesize that some of these *Os*ANN proteins may participate in rice-BPH interactions. To test this hypothesis, we first detected the transcript levels of *OsANN* genes before and after BPH feeding. Expression of most *OsANN* genes was induced by BPH feeding, with the expression levels of *OsANN2*, *OsANN*6, *OsANN*8 being the highest (*SI Appendix*, Fig. S10). Next, we produced stable transgenic Nipponbare plants overexpressing *OsANN2*, *OsANN6* or *OsANN8*. We were able to obtain *OsANN2* and *OsANN8*, but not *OsANN6* transgenic lines. Accordingly, several independent lines of *OsANN2*- and *OsANN8*-expressing transgenic plants (*SI Appendix*, Fig. S11*A* and Fig. S11*B*) were further analyzed.

There were no significant growth differences between wild-type Nipponbare and *OsANN2* and *OsANN8* overexpressing plants (*SI Appendix*, Fig. S11*C*–*E* and S12). We evaluated the survival rates of BPH feeding on the *OsANN*-expressing plants. *OsANN2*-OE and *OsANN8*-OE rice plants displayed enhanced resistance against BPH as the survival rate of insects were markedly decreased when fed on *OsANN2*-OE and *OsANN8*-OE rice plants compared to that on Nipponbare plants (Fig. 5*A* and 5*B*). Consistent with this observation, the survival rates of *OsANN2*-OE and *OsANN8*-OE rice plants infested by ds*GFP* BPH were higher than Nipponbare plants (*SI Appendix*, Fig. S5*C* and S5*D*). We also measured the honeydew production of *dsGFP* BPH that fed on Nipponbare, *OsANN2*-expressing and *OsANN8*-expressing rice plants (*SI Appendix*, Fig. S5*F* and S5*G*). ds*GFP* BPH fed on *OsANN2*-OE and *OsANN8*-OE rice seedlings had mostly lower amounts of excreted honeydew compare to those fed on Nipponbare plants.

**Fig. 5.**
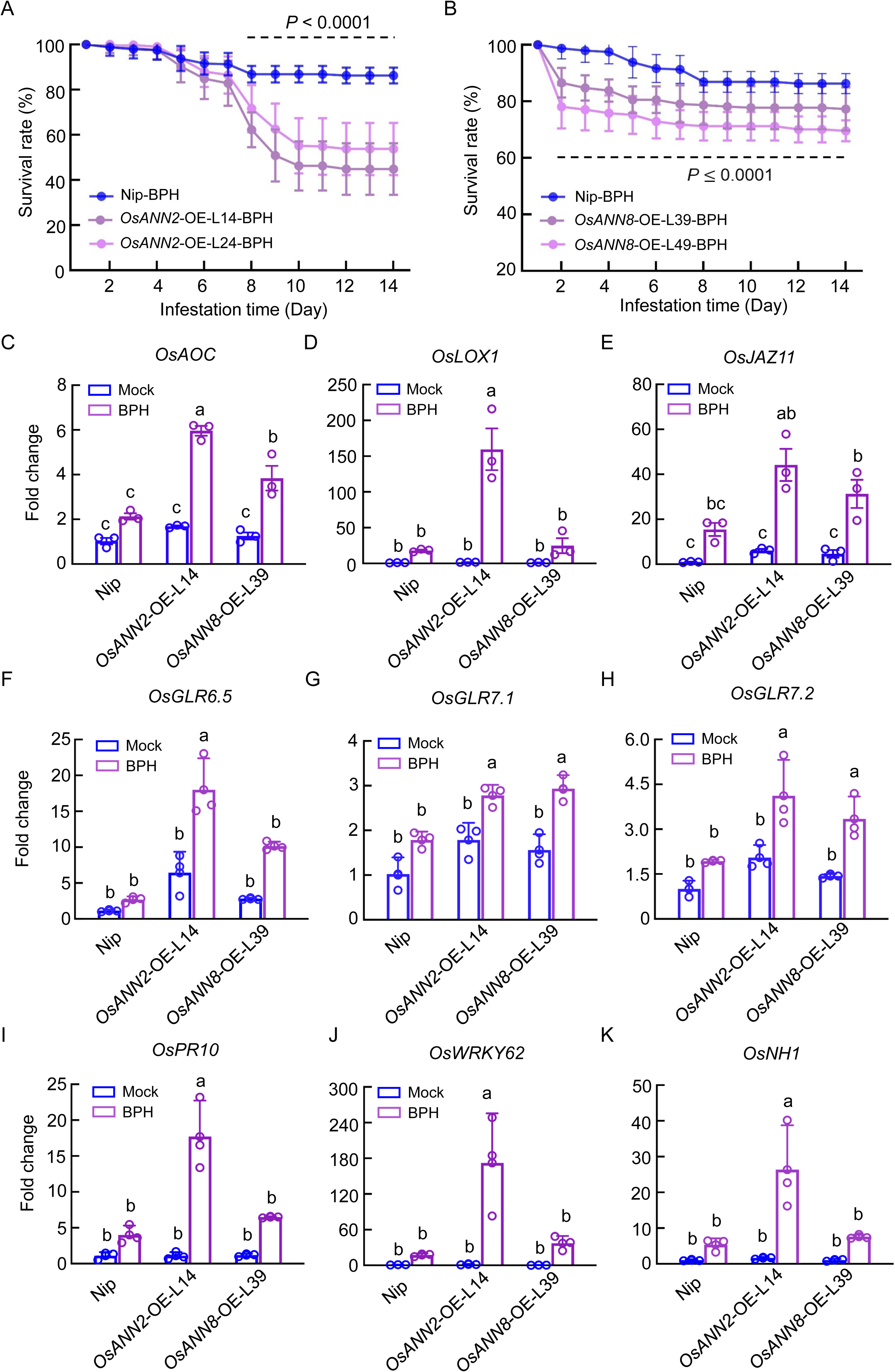
*Os*ANN2 and *Os*ANN8 positively regulate rice resistance against BPH. (*A* and *B*) The daily survival rates of wild type BPH fed on 5-leaf-stage of Nip, *OsANN2*-OE and *OsANN8*-OE plants. Values are represented as mean ± s.e.m. of 8 biological replicates (Two-way ANOVA; 20 individual 2^nd^ BPH nymphs per rice plant for each biological replicate). (C*–E*) Transcript levels of JA biosynthesis genes *OsAOC* and *OsLOX1* as well as JA responsive gene *OsJAZ9* in Nip, *OsANN2*-OE and *OsANN8*-OE plants fed by 20 individual 5^th^ instar BPH for 24 h. (*F–H*) Transcript levels of *OsGLR* genes in Nip, *OsANN2*-OE and *OsANN8*-OE plants fed by 20 individual 5^th^ instar BPH for 24 h. (*I–K*) Transcript levels of SA relevant genes, *OsPR10*, *OswRKY62* and *OsNH1*, in Nip and *OsANN2*-OE plants fed by 20 individual 5^th^ instar BPH for 24 h. (*L–N*) Transcript levels of SA pathway genes, *OsPR10*, *OswRKY62* and *OsNH1,* in Nip and *OsANN8*-OE plants fed by 20 individual 5^th^ instar BPH for 24 h. Values are displayed as mean ± s.e.m. (*n* = 3 biological replicates) and different letters indicate statistically significant differences analyzed by two-way ANOVA (Tukey test, *P* < 0.05) in (*C–N*). The *OsACTIN* (*Os03g50885*) gene was used as internal control. Experiments were repeated three times.

Given that *Nl*ANX5 facilitates the survival of BPH on Nipponbare plants and down-regulates JA biosynthesis and signaling pathway genes, we next examined JA genes expression in *OsANN2*-OE and *OsANN8*-OE plants. Interestingly, *OsAOC*, *OsLOX1*, *OsJAR2, OsJAMYB*, *OsJAZ9* and *OsJAZ11* were all up-regulated at higher levels in *OsANN2*-OE and *OsANN8*-OE rice plants than in Nipponbare plants, especially in *OsANN2*-OE plants (Fig. 5*C*–*E*; *SI Appendix*, Fig. S13*A*–*C*). In addition, the transcript levels of *OsGLR6.5*, *OsGLR7.1* and *OsGLR7.2* were higher in *OsANN2*-OE and *OsANN8*-OE plants when compared to that in Nipponbare plants (Fig. 5*F*–*H*). Finally, expression of SA pathway marker genes (48−51), such as *OsPR10*, *OsWRKY62*, *OsPAL1*, *OsNH1* and *OsNH2*, were augmented in both *OsANN2*-OE and *OsANN8*-OE plants in response to BPH feeding (Fig. 5*I*–*K*; *SI Appendix*, Fig. S13*D*–*E*). Taken together these results suggest that *OsANN2* and *OsANN8* positively regulate multiple rice defenses against BPH attacks through augmenting GLR, JA and SA pathway gene expression.

### Further characterization of *Os*ANN-mediated rice resistance against BPH

To further investigate the relationship between BPH *Nl*ANX5 and rice *Os*ANN2 and *Os*ANN8, we constructed the *OsANN2* and *OsANN8* double knockout mutant, *osann2 osann8*, in the genetic background of Nipponbare (Fig. 6*A*). Nipponbare and *osann2 osann8* plants were then fed with ds*GFP* BPH or ds*NlANX5* BPH. As shown from Fig. 6*B*, ds*NlANX5* BPH displayed a lower survival rate on wild-type Nipponbare plants when compared with ds*GFP*, consistent with the previous results (Fig. 2*C*, 5*A* and 5*B*). Remarkably, loss-of-function of *OsANN2* and *OsANN8* almost completely restored the survival of ds*NlANX5* BPH (Fig. 6*B*). More honeydew was produced by BPH insects in the *osann2 osann8* rice seedlings than in Nipponbare plants (*SI Appendix*, Fig. S5*H*) and, conversely, fewer *osann2 osann8* rice seedling survived infestation by ds*NlANX5* BPH (*SI Appendix*, Fig. S5*B*). We were also successful in generating the *osann2* single mutant rice plants (*osann2*-1 and *osann2*-2). As shown in *SI Appendix*, Fig. S14, the *osann2* single mutant rice plants (*osann2*-1 and *osann2*-2) could not fully restore the survive of ds*NlANX5* BPH, suggesting that *Os*ANN2 and *Os*ANN8 work additively or in concert to resist BPH in rice plants. Together, these results provide strong genetic evidence that *Os*ANN2 and *Os*ANN8 are major molecular targets of *NI*ANX5 for BPH survive of BPH on rice.

**Fig. 6.**
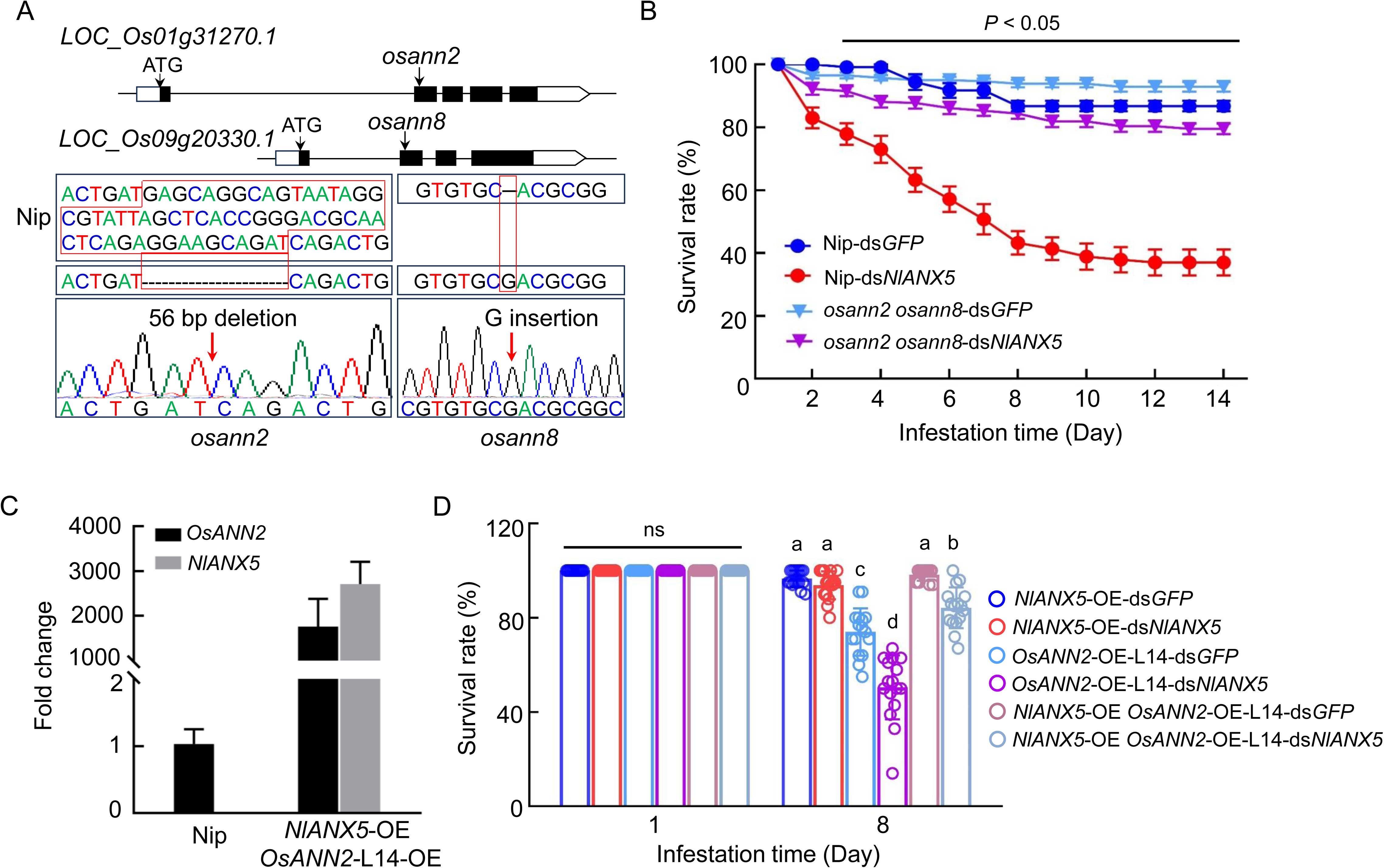
*Nl*ANX5 inhibits *Os*ANN-mediated rice resistance against BPH. (*A*) Mutations generated in *OsANN2* and *OsANN8* by CRISPR/Cas9. Introns and exons are shown as lines and boxes, respectively. Black box, translated region; white box, untranslated region; ATG, translation start site. The mutated sites in *osann2* and *osann8* are indicated. (*B*) The daily survival rates of ds*GFP* and ds*NlANX5* BPH insects fed on Nip and *osann2 osann8* double mutant plants. Values are represented as mean ± s.e.m. of 8 biological replicates (Two-way ANOVA; 20 individual 2^nd^ BPH nymphs per rice plant for each biological replicate). (*C*) Transcript levels of *NIANX5 and OsANN2* in *NlANX5*-OE *OsANN2*-OE double overexpressing rice plants. Nip was used as a control. Values are displayed as mean ± s.e.m. (*n* = 3 biological replicates). (*D*) The survival rates of ds*GFP* and ds*NlANX5* BPH insects fed on Nip, *OsANN2*-OE*, NlANX5*-OE *and OsANN2 NlANX5*-OE plants at day 1 and day 8. Values are represented as mean ± s.e.m. of 8 biological replicates (Two-way ANOVA; 20 individual 2^nd^ BPH nymphs per rice plant for each biological replicate). Experiments were repeated three times.

We next constructed the *NlANX5*-OE *OsANN2*-OE double overexpressing plants in the background of Nipponbare (Fig. 6*C*) to further investigate the opposing functions of BPH *Nl*ANX5 and rice *Os*ANN2 during BPH infestation. Notably, the ability of *OsANN2*-OE to counter the virulence function of *Nl*ANX5 is reduced when *Nl*ANX5 is transgenically overexpressed (i.e., in *OsANN2 NlANX5*-OE plants; Fig. 6*D*). This result suggests that, while transgenic overexpression of *OsANN2* can effectively counter *Nl*ANX5 naturally delivered into rice cells, which is presumably at a low amount, during BPH infestation (Fig. *5A*), the relative amounts of *Os*ANN2 and *Nl*ANX5 determine the outcome of the opposing functions of *Nl*ANX5 and *Os*ANN2, which provide further evidence for a competition-based mechanism between BPH *NI*ANX5 and rice *Os*ANN2 proteins during the BPH-rice interaction.

## Discussion

In this study, we show that *Nl*ANX5 is a crucial BPH virulence effector delivered to the rice plant to target and counter host annexin proteins which we found to play a major role in rice resistance to BPH. We observed that *NlANX5*-RNAi BPH has a longer non-penetration phase but a shortened phloem sap uptake phase (Fig. 2*D*–2*F*), which is correlated with an eventual lower survival rate on Nipponbare plants compared to control *GFP*-RNAi BPH (Fig. 2*C*). Our observations that *NIANX5* is expressed in the salivary glands of BPH (Fig. 1*A*) and detected in rice sheath fed by BPH (Fig. 1*B*) and that the feeding and survival defects of *NlANX5*-RNAi BPH insects could be restored in *NlANX5*-expressing rice plants (Fig. 2*C*) are all consistent with the hypothesis that *Nl*ANX5 functions in rice tissues. We identified rice annexins as the host targets of *Nl*ANX5 and showed that *Nl*ANX5 suppresses the rapid [Ca^2+^]_cyt_ influx and expression of multiple defense pathway genes (Fig. 3). Overall, this study not only provides a previously uncharacterized mechanism by which an insect annexin targets host annexin-mediated Ca^2+^ and associated defense pathways to facilitate insect feeding and survival in host plants, but also illustrates an example of using the host targets, *Os*ANN2 and *Os*ANN8, as a promising source of pest resistance.

In plants, Ca^2+^ influx in cytoplasm is an early and crucial step against pathogen attacks (37, 52−53). Plants have evolved diverse membrane Ca^2+^ channels to facilitate increases in cytosolic Ca^2+^ concentrations. Some calcium channels, such as OSCA1.3, cyclic nucleotide-gated channel (CNGC), WeiTsing and resistosomes, gate plant immune responses against pathogen by triggering influx of calcium ions across the plasm membrane (10, 12, 54−55). In addition, other Ca^2+^ transporters, including a vacuolar Ca^2+^/H^+^ exchangers (CAXs), can be activated to scavenge excess Ca^2+^ for balancing plant growth and immunity (56). These studies have greatly broadened our understanding of the crucial roles of Ca^2+^ in plant-pathogen interactions. However, our knowledge of Ca^2+^ dynamics in the co-evolution of plants and insects are still limited. In the present study, we found that *Os*ANN2 and *Os*ANN8 positively regulate rice resistance against BPH (Fig. 5*A* and 5*B*), and this process is associated with the boosted expression of *GLRs* and JA and SA pathway genes (Fig. 5*C*–*K*). *Os*ANN genes are themselves induced during BPH feeding in rice (*SI Appendix*, Fig. S10). Thus, *Os*ANNs are likely part of a positive feedforward mechanism for *GLR*-mediated Ca^2+^ spikes and JA and SA defense activation during BPH feeding. Our results suggest that this Ca^2+^ feedforward mechanism is subverted by a major BPH virulence effector, *Nl*ANX5.

How *Os*ANN2 and *Os*ANN8 boost Ca^2+^ response during BPH feeding represents an important question to be elucidated in future studies. Annexins are Ca^2+^- and charged phospholipid-binding proteins (20, 57−58). Plants annexins were suggested to allow for influx of cytoplasmic Ca^2+^ during stress responses. A *capsicum* annexin had Ca^2+^ channel activity *in vitro* and might affect Ca^2+^ influx in plant cells (59). Maize (*Zea mays*) annexins exhibit active Ca^2+^ conductance in lipid bilayers and were also displayed to exhibit channel activity in oxidized membranes (60−61). Root cell-expressed *AnnAt1* was also characterized to exhibit channel activity (62). However, whether *Os*ANN2 and *Os*ANN8 form Ca^2+^-channels to directly transmit Ca^2+^ is not known. In this study, we found that *Os*ANN2 and *Os*ANN8 can up-regulate the expression of *GLR* genes (Fig. 5*F*–*H*), suggesting they may augment GLR channel activity through enhanced *GLR* gene expression to promote Ca^2+^ influx in response to BPH feeding. However, since annexin proteins are themselves associated with Ca^2+^ and phospholipids in the membrane (20, 58, 63−64), it is possible that the enhanced *GLR* gene expression is a result of a more direct impact of *Os*ANN proteins on the Ca^2+^ permeability of the GLR channels in the membrane.

It is notable that *Nl*ANX5 can displace *Os*ANN2 and *Os*ANN8 from the membrane (Fig. 4*E*and 4*F*), suggesting a direct competition mechanism. To our knowledge, this may be the first example in which an insect annexin has evolved to interfere with the membrane association of the host annexins to exert its virulence function. Future advanced cellular imaging methodologies are needed to gain further insights into how *Nl*ANN5 displaces *Os*ANN2 at the rice cell membrane. In addition, future research should examine the sequence features of *Nl*ANX5 and *Os*ANNs that confer the apparently opposing functions during the rice-BPH interaction. *Nl*ANX5 contains two annexin domains (25), whereas *Os*ANN2, *Os*ANN8 and other *Os*ANNs have three or more annexin domains (58). There are also other sequence differences between *Nl*ANN5 and *Os*ANNs (*SI Appendix* Fig. S15).

While our study focuses on *Nl*ANX5 in BPH, we note that annexin proteins are found in diverse insects and plants (*SI Appendix* Fig. S16). In addition, a putative annexin-like effector is present in the secreted saliva of white-back planthopper (*Sogatella furcifera*) (8) and the cereal cyst nematode (*Heterodera avenae*) (65). The host targets of these insect/nematode annexins remain unknown. Our identification of host annexins as the molecular targets of *Nl*ANX5 in the rice-BPH interaction raises the possibility that annexin-based molecular battles between plants and insects/nematodes may be common. If so, enhanced expression of host annexins may be explored as a broadly applicable anti-virulence breeding strategy against diverse insect and nematode pests.

## Materials and Methods

Experimental materials and methods, including plant and insect materials, transgenic rice production, *in situ* mRNA hybridization analysis, rice sheath sample preparation for LC-MS analysis, RNA interference, BPH survival test on rice, Ca^2+^ imaging and analyses, electrical penetration graph, co-immunoprecipitation assays, bimolecular fluorescence complementation assays, protoplast isolation and transfection, protein extraction and western blot, RNA isolation and quantitative real-time PCR, and AlphaFold3 predictions can be found in *SI Appendix, Supplementary Materials and Methods*.

## Data Availability and Sharing Plan

All data are presented within this paper and will be published open access. All biomaterials will be available to researchers without restrictions

## Acknowledgments

This study was supported by fundings from the National Key Research and Development Plan in the 14^th^ five years plan (Grant 2021YFD1401100 to C.X.Z.), the Yunnan Fundamental Research Projects (Grants 202401AS070122 and 202505AS350005 to Y.J.J.; Grant 202105AC160082 to Y.J.J., which supported her research at Michigan State University), Zhejiang University Student International Exchange Fund to X.Y.Z., which supported her research at Michigan State University), and Howard Hughes Medical Institute to S.Y.H. We thank to the Plant Molecular Biology Research Team of Yunnan University for experimental platform support to Y.J.J. during this work.

