## Supplemental Information for "Augment rice *ANNEXIN* expression to counter planthopper *Nl*Annexin-like5 as an anti-virulence strategy against a major crop pest"

PNAS
www.pnas.org

SUPPLEMENTARY INFORMATION

### Augment rice *ANNEXIN* expression to counter planthopper *Nl*Annexin-like5 as an anti-virulence strategy against a major crop pest

Xiao-Ya Zhang [a,b], Shaoqin Li[c,d], Comzit Opachaloemphan[g], Chuan-Xi Zhang [a,e,*], Sheng Yang He[b,f,g,*], Yanjuan Jiang [b,c,d,*]

[a] Institute of Insect Science, Zhejiang University, Hangzhou, 310058, China.

[b] DOE Plant Research Laboratory, Michigan State University, East Lansing, MI, USA.

[c] State Key Laboratory for Conservation and Utilization of Bio-Resources in Yunnan, School of Life Sciences, Yunnan University, Kunming 650091, China.

[d] CAS Key Laboratory of Tropical Plant Resources and Sustainable Use, Xishuangbanna Tropical Botanical Garden, Chinese Academy of Sciences, Kunming, Yunnan 650223, China.

[e] State Key Laboratory for Quality and Safety of Agro-Products, Institute of Plant Virology, Ningbo University, Ningbo 315211, China.

[f] Howard Hughes Medical Institute, Duke University, Durham, NC, USA.

[g] Department of Biology, Duke University, Durham, NC, USA.

*Corresponding authors

**This PDF file includes:**

Supplementary text (SI Materials and Methods)

Figures S1 to S17

Tables S1 to S3

SI References

#### SI Materials and Methods

**Plant materials and insects.** *Oryza sativa* ssp. Japonica cv. Nipponbare was used as the wild type control for most experiments and for generating transgenic plants. The *Nilaparvata lugens* population used in this study was created artificially over many generations beginning in 2008 from a wild rice field on the Huajiachi Campus of Zhejiang University in Hangzhou, China (30° 16' N, 120° 11' E). BPH insects were raised in environmentally controlled growth chambers on the susceptible cultivar rice variety Xiushui 134, which are reared in a modified Yoshida solution with a pH of 4 (in which ammonium nitrate was substituted by 1.06 M carbamide), at a temperature of 27 ± 0.5°C and a relative humidity of 50 ± 0.5% and a photoperiod of 16 h of light and 8 h of darkness.

**Transgenic rice.** To create the 35S:*Nl*ANX5-YFP construct, the *NlANX5* coding sequence was amplified from BPH and recombined into the intermediate vector pDonor207 using the BP recombination kit (Invitrogen) and then into the destination vector pEarleygate104 using the LR recombination kit (Invitrogen). Similarly, the *OsANN2* (*LOC_Os01g31270*) and *OsANN8* (*LOC_Os09g20330*) coding sequences were amplified from rice (*Oryza sativa* subsp. japonica) cultivar Nipponbare and inserted into the destination vector pEarleygate101. The transgenic rice plants were created using Agrobacterium-mediated transformation at BIOGLE GeneTech (Hangzhou Biogle Co., LTD.), where rice plants were grown in a greenhouse at 27 ± 0.5°C and 60 ± 0.5% relative humidity with a 16-hour light and 8-hour dark photoperiod.

For *OsANN2* gene editing, a single target site was inserted between *OsU6a* and sgRNA in the pYLsgRNA-*OsU6a* vector (1), and the *OsU6a*-sgRNA cassette was subsequently cloned into the pMH-SA vector using the restriction enzyme sites *Spe*I and *Asc*I (2). A tandem combination of *OsU6a*-sgRNA (for *OsANN2*) and *OsU6b*-sgRNA (for *OsANN8*) was cloned between *Asc*I and *Spe*I in the pMH-SA vector for double editing of *OsANN2* and *OsANN8*. Sequencing was used to identify homozygous mutant lines. Supplementary Table S2 lists the primers used in vector production and genome editing.

***In situ* mRNA hybridization analysis.** Fresh salivary tissues were isolated from 100 5$^{th}$ instar BPH and washed 3 times before being placed in the 4% paraformaldehyde (PFA) fixation solution (Servicebio, G1113) at 4°C overnight, dehydrated and embedded in paraffin. Ten μM-thick sections were cut on glass slides, dewaxed and digested by Protease K (Servicebio, G1205) at 37°C for 15 min. After three washes in phosphate-buffered saline (PBS) solution (Servicebio, G0020), the sections were incubated in pre-hybridization solution at 37°C for 1 h and then in hybridization solution containing the *NlANX5* probe (1 μM) at 42°C overnight. The *NlANX5* probe (5'-CY3-CAATA GTGGC AGCC A TCAGC TCAGG CATAG-CY3-3') has double-ended marking CY3 tag generated using the DIG RNA Labeling Kit (Roche Applied Science, Penzberg, Germany) based on the manufacturer's instructions. The slides were stained with DAPI for 8 min in the dark after being washed in saline sodium citrate (SSC) buffer (Servicebio, G3016-4) for three times (2 × SSC, 10 min; 1 × SSC, 2 × 5 min; 0.5 × SSC, 10 min). Sections hybridized with sense probe were used as a negative control and antisense probe was used to detect the target mRNA. All steps were performed at 37°C and sections were examined under a Nikon fluorescence microscope with a 40 × lens (330 nm–380 nm for DAPI, 510 nm–560 nm for CY3).

**Rice sheath and phloem exudates for LC-MS.** Nine independent sets of Nipponbare rice sheath protein samples (three biological replicates) were collected after feeding by 5$^{th}$ instar nymphs as follows. Nine 6-leaf-stage rice plants were individually placed into a breathable transparent plastic bottle (5 cm in diameter, 30 cm in height) containing Yoshida medium, pH = 4, and then fed by ~500 fifth instar BPH for 72 h. Next, the two outermost layers of leaf sheath surrounding the stem were cut into small pieces, rinsed with ultrapure water for three times and ground to powder in liquid nitrogen. Total protein was extracted in a buffer containing 60 mM Tris-HCl (pH = 7.6), 2.5% Glycerol, 6% SDS, 0.1 M DL-Dithiothreitol (DTT), 1x Protein inhibitor cocktail and 1 mM phenylmethanesulfonyl fluoride. Protein samples were concentrated using 3-kDa filtration units and shipped on dry ice to Shanghai Hoogen Biotech for shot-gun LC-MS analysis. Briefly, protein samples digested by trypsin for 16 h at 37°C. The hydrolysates were then loaded into Zorbax 300SB-C18 peptide traps (Agilent Technologies, Wilmington, DE) and analyzed by a Q Exactive mass spectrometer (Thermo Fisher). The raw LC-MS data were searched against the BPH protein database (NCBI Accession No. PRJNA669454) using software MaxQuant 1.5.5.1. Peptides with a high confidence score was extracted, and the database search was conducted with false discovery rate (FDR) under 0.01.

Phloem exudate collection was performed on four-leaf-stage rice seedlings following an optimized EDTA-enhanced exudation protocol adapted from established methodologies (3,4). The experimental procedure involved excising plant shoots approximately 1 cm above the root-shoot junction using sterile razor blades. To eliminate contaminants from wound-induced root exudation, the decapitated shoots were initially immersed in 20 mM EDTA buffer (pH 8.0, adjusted with KOH) and vortexed for 15 seconds. Subsequently, 4-6 prepared shoots were incubated in sterile polypropylene tubes containing 1.5 mL fresh EDTA solution (20 mM, pH 8.0) under controlled environmental conditions (darkness, >90% humidity) for 12 h to minimize transpirational water loss. Post-incubation, pooled exudates from 40 biological replicates underwent centrifugal concentration using 10 kDa molecular weight cutoff filters at 4°C. The final concentrated extracts (20 μL total volume) were aliquoted for subsequent mass spectrometry analyses.

**RNA interference (RNAi).** The nucleotide sequence (around 500 bp long) specific to the *NlANX5* coding region was cloned into the pMD 19-T vector (TaKaRa). The double-stranded RNA was synthesized through PCR amplification by using the Mega script T7 High Yield RNA Transcription Kit (Vazyme, Nanjing, China), following the manufacturer's instructions. The procedure for gene knockdown by RNAi was described previously. Briefly, the first and fifth instar BPH nymphs in an insect rearing growth chamber were anaesthetized with carbon dioxide for ~10 s. Approximately 30-250 ng of dsRNA was injected into BPH mesothorax using FemtoJet (Eppendorf-Netheler-Hinz, Hamburg, Germany). The efficiency of RNA interference was assessed 48 h after injection. RNA from 20 nymphs was extracted using the RNAiso Plus kit (TaKaRa) according to the manufacturer's instructions as one biological replicate. The relative level of the *NlANX5* transcript was quantified using a CFX96TM Real-Time PCR Detection System (Bio-Rad, Hercules, CA, USA) with primers shown in Supplementary Table S2. The Aequorea Victoria green fluorescent protein (GFP) gene sequence was used as the control template in RNAi experiments. Three biological replicates were conducted, and each has three technical replicates.

**BPH survival test on rice.** BPH nymphs with verified *NlANX5* knockdown by RNAi were placed into a small breathable transparent plastic bottle of 10 cm in diameter and 9 cm in height containing one-week-old rice seedlings (1-leaf-stage) for 24 h to recover. About 20 healthy-appearing nymphs were gently transferred into a longer breathable transparent plastic bottle of 5 cm in diameter and 30 cm in height containing four-week-old rice plants (4- to 5-leaf-stage) and observed for up to 15 days. The feeding space for BPH in each plastic bottle was around 200 $cm^3$ divided by two sponges of 5 cm in diameter and 2 cm in thickness. The number of surviving BPHs were counted daily on each plant. The ds*GFP* injected BPH were used as control. The average survival rate on each seedling was calculated.

**Cyto-$Ca^{2+}$ imaging and analyses.** A ratiometric calcium indicator, RGECO1-mTurquoise (5), was introduced into Nipponbare plants to generate the cyto-$Ca^{2+}$-sensor. A total of twelve independent transgenic lines were produced and six lines were showed to overexpression both RFP and mTurquoise transcripts. These RGECO1-mTurquoise-expressing plants were propagated to T3 generation and line 8 was selected for further $Ca^{2+}$ assay. Plants of line 8 displayed no noticeable growth defects compared to Nipponbare plants (*SI Appendix*, Figure S4). Plants with the R-GECO1 cyto-calcium sensor were grown in the modified Yoshida solution for 4 weeks. Microscopic analyses were performed using an In Vivo Imaging System (IVIS) (PerkinElmer, Waltham, USA), which uses the optical imaging technology to facilitate non-invasive longitudinal monitoring of cells with twenty-eight high efficiency filters spanning 430–850 nm. The R-GECO1 signal was imaged with $\lambda_{Ex}$ = 532 nm and $\lambda_{Em}$ = 580–640 nm, whereas mTurquoise signal was imaged with $\lambda_{Ex}$ = 405 nm and $\lambda_{Em}$ = 460–520 nm. The CCD camera was cooled to −90°C before use and the chamber temperature was set to 27°C. Rice plants were transferred into a glass tube (2 cm in diameter, 15 cm in length) with 30 5th instar BPH nymphs each. The glass tubes containing plants and BPHs were fixed on the imaging chamber shelf with a black electrical tape and imaged. The area with BPH feeding (2 cm in length) was used for ROI (Region of interest) calculation and plants without BPH feeding were used as control. After background subtractions ratiometric image calculations followed that described by Kardash *et al.* (6). R-GECO1-mTurquoise ratio changes (ΔR: R) were calculated from background-corrected images and normalized to the average of the initial baseline (five frames at the start of the measurement, 1–5 min). T Statistics were performed using Living Image® Version 4.5 Software (PerkinElmer, Waltham, USA) and $n \geq 6$ plants per treatment. Free $[Ca^{2+}]_{cyt}$ from ΔR:R values were calculated according to:

$$[\mathrm{Ca}^{2+}]_{cyt} = K_d \times (\frac{\Delta R/R \quad - \Delta R/R_{min}}{\Delta R/R_{max} - \Delta R/R})^{1/n}$$

where $\Delta R:R_{min}$ is the minimum ratio change, $\Delta R:R_{max}$ is the maximum ratio change, n represents the Hill coefficient (1.84) and Kd represents the apparent dissociation constant for $Ca^{2+}$ determined for R-GECO1-mTurquoise (149 nM; Supplementary Table S2).

**Electrical penetration graph (EPG).** A Giga-8 DC EPG amplifier (Wageningen Agricultural University, The Netherlands) was used to record the different feeding behaviors of BPH on rice plants (7–9). The early 5th instar BPH nymphs were firstly treated with ds*NlANX5* or ds*GFP* and fed on 1-week-old Nipponbare rice plants. After being reared for three days, the adult BPHs were carefully transferred by a soft bristled pen, without being anesthetized by $CO_2$, and their thoraxes were attached to a gold wire (12 μm in diameter, 30 mm in length) with a water-soluble conductive silver glue

(Wageningen Agricultural University, The Netherlands) under an optical microscope. The end of the gold wire was connected to the amplifier through an EPG probe and a copper electrode (2 mm in diameter, 10 cm in length) was inserted into the nutrient solution of rice plants to establish the other part of the electrical circuit. The EPG recording was performed in a climate-controlled room (26 ± 1°C) for 20 h including 8 h under light and 12 h in the dark continuously. EPG data were analyzed using EPG Stylet+ software (Wageningen Agricultural University, The Netherlands) and each treatment had at least ten biological replications. There are five classic waveforms, which can be distinguished by EPG: (1) NP, the non-penetration phase; (2) PP, the pathway phase including extracellular activities before, after or between N1–3, during which the wave form is not as regular as N1–3 phases; (3) N1–3, including N1 penetration initiation, N2 salivation and stylet movement and N3 extracellular activity near the phloem; (4) N4, the intracellular phase of activity in the phloem and the phloem sap ingestion phase; (5) N5, the intracellular phase of activity in the xylem and the phloem sap ingestion phase (5–7).

**Co-immunoprecipitation (IP) assays**. The full-length cDNA of *NlANX5* was cloned into the pEG104 destination vector to produce a fusion with YFP (35S:YFP-*Nl*ANX5). The full-length cDNAs of *OsANNs* were inserted into the vector pGWB517 to obtain in-frame fusions with Myc (35S:Os*ANN*s-Myc). the 35S:YFP construct was used as a negative control. The resulting plasmids were transformed into *Agrobacterium tumefaciens* (*A. tumefaciens*) strain GV3101, and different pairs of *A. tumefaciens* transformants were then infiltrated into *Nicotiana benthamiana* (*N. benthamiana*) leaves by hand. Forty-eight hours after infiltration, 1g leaf samples for each treatment were collected to perform the co-IP assay. Total protein was extracted with a co-IP buffer containing 50 mM Tris-HCl [pH 7.5], 150 mM NaCl, 0.2% v/v Triton X-100, 50 μM MG132, 5 mM dithiothreitol, and 1x protease inhibitor cocktail. Immunoprecipitation experiments were performed with GFP-trap beads following the manufacturer's protocol. In brief, cell lysates were incubated with GFP-trap beads (catalog no. gta-100, ChromoTek, Munich, Germany) for 4 h at 4°C. After incubation, the beads were washed four times with the extraction buffer and the co-immunoprecipitated proteins were then detected by immunoblotting using an anti-Myc antibody (catalog no. ab9106, Abcam, Cambridge, UK; 1:10,000). The YFP-*Nl*ANX5 fused protein was detected using an anti-GFP antibody (catalog no. ab290, Abcam, Cambridge, UK; 1:10,000). Experiments were repeated at least four times. The primers used for the vector construction are listed in Supplemental Table S2.

**Bimolecular fluorescence complementation (BiFC) assays.** The cDNA sequences encoding the 64 amino acids of the C-terminal end of enhanced YFP (cYFP) and the 173 amino acids of the N-terminal end of YFP (nYFP) were PCR-amplified and individually inserted into pFGC5941 plasmids to produce pFGC-cYFP and pFGC-nYFP, respectively (10). The full-length cDNA of *NlANX5* was cloned into pFGC-cYFP to produce fusion with cYFP (*Nl*ANX5-cYFP). The full-length sequences of *Os*ANN2, *Os*ANN6 and *Os*ANN8 were inserted into pFGC-nYFP to generate in-frame fusions with nYFP (*Os*ANN2-nYFP, *Os*ANN6-nYFP and *Os*ANN8-nYFP), respectively. The resulting plasmids were transformed into *A. tumefaciens* strain GV3101, and different pairs of *A. tumefaciens* transformants were then infiltrated into *N. benthamiana* leaves by hand. Forty-eight h after infiltration, YFP fluorescence was examined under a confocal laser-scanning microscope (Olympus, Tokyo, Japan). YFP was detected using white light laser (WLL) at 40% intensity, and Photomultiplier Module (PMT) at 65% gain with excitation wavelength of 514 nm and emission at 524–570 nm. The experiments were performed at least four times using different batches of *N. benthamiana* plants. The primers used for vector construction are listed in Supplemental Table S2.

**Protoplast isolation and transfection.** Isolation of rice protoplast was carried out as described (11). PEG-mediated transfection was used with some modifications. Briefly, for each samples (*Os*ANN2-HA, *Os*ANN6-Myc, *Os*ANN8-Myc as well as *Os*ANN2-HA + *Nl*ANX5-Flag, *Os*ANN6-Myc + *Nl*ANX5-Flag and *Os*ANN8-Myc + *Nl*ANX5-Flag), around 10–30 µg of plasmid DNA were mixed with 500 µL protoplasts (about 5 x $10^5$ cells). 550 µL freshly prepared PEG solution [40% (W/V) PEG4000, 0.6 M mannitol and 0.1 M $CaCl_2$] were added. The mixture was incubated at room temperature for 20 min. After incubation, 2.2 mL W5 solution (154 mM NaCl, 125 mM $CaCl_2$, 5 mM KCl and 2 mM MES at pH 5.7) were added slowly, then mixed well by gently inverting the tube. The protoplasts were pelleted by centrifugation at 200 g for 2 min, then resuspended gently in 5 mL W5 solution for 16 h culturing at 28°C. Samples (*Os*ANN2-Myc + YFP and *Os*ANN2-Myc + *Nl*ANX5-YFP, *Os*ANN6-Myc + YFP and *Os*ANN6-Myc + *Nl*ANX5-YFP as well as *Os*ANN8-Myc + YFP and *Os*ANN8-Myc + *Nl*ANX5-YFP) were transfected for subsequent co-IP assay in the presence of $Ca^{2+}$.

**Protein extraction and western blot.** 0.5 mL transfected protoplasts was used for total protein extraction, and the rest of protoplasts (4.5 mL) was used for protein extraction from soluble vs. membrane fractions. Protoplasts were harvested by centrifugation at 300 g for 3 min. Total protein was extracted with 100 µL protein buffer (0.33 M sucrose, 0.05 M Mops, 5 mM EDTA, 5 mM $CaCl_2$, 2 mM DTT, 1 mM PMSF, 0.5% Triton and 1x protein inhibition cocktail at pH 6.8). Soluble protein was extracted with 900 µL soluble protein extraction buffer (0.33 M sucrose, 0.05 M Mops, 5 mM EDTA, 10 mM $CaCl_2$, 2 mM DTT, 1 mM PMSF and 1 x protein inhibition cocktail at pH 6.8), while the pellet was extracted with 900 µL membrane protein extraction buffer (0.33 M sucrose, 0.05 M Mops, 5 mM EDTA, 10 mM $CaCl_2$, 2 mM DTT, 1 mM PMSF, 0.5% Triton and 1 x protein inhibition cocktail at pH 6.8). After extraction, half of the soluble protein or membrane protein fraction were subjected to immunoprecipitation with HA-trap beads (catalog no. M20013S, Abmart, Shanghai, China), and the other half were immunoprecipitated with Flag-trap beads (catalog no. PFA050, LABLEAD, Beijing, China). Equal volumes of total, soluble protein and membrane protein were used for western blot to detect *Os*ANN2-HA, *Os*ANN8-Myc or *Nl*ANX5-Flag proteins. *Os*ANN2-HA protein was detected by using an anti-HA antibody (catalog no. M20003, Abmart, Shanghai, China; 1:5,000), *Os*ANN8-Myc protein was detected by using an anti-Myc antibody (catalog no. M20002, Abmart, Shanghai, China; 1:5,000) and *Nl*ANX5-Flag protein was detected by using an anti-Flag antibody (catalog no. M20008, Abmart, Shanghai, China; 1:5,000).

For co-IP assay in the presence of $Ca^{2+}$, protoplasts expressing GFP, *Nl*ANX5-GFP with *Os*ANN2-Myc and *Os*ANN8-Myc were harvested by centrifugation at 300 g for 3 min. Total protein was extracted with 500 µL protein buffer (0.33 M sucrose, 0.05 M Mops, 5 mM EDTA, 2 mM DTT, 1 mM PMSF, 0.3% Triton and 1x protein inhibition cocktail at pH 6.8). After extraction, 100 µL were separated from each sample as the input. The remaining extraction was divided into two equal parts and fresh protein buffer was added to each part to 1 mL, and an additional 5 mM of $CaCl_2$ was added to one of the parts. Then each samples were subjected to immunoprecipitation with GFP-trap beads (catalog no. gta-100, ChromoTek, Munich, Germany). Equal volumes of input, IP with $Ca^{2+}$ and IP without $Ca^{2+}$ were used for western blot to detect *Os*ANN2-Myc, *Os*ANN8-Myc or *Nl*ANX5-YFP proteins. *Os*ANN2-Myc or *Os*ANN8-Myc protein was detected by using an anti-Myc antibody (catalog no. M20002, Abmart, Shanghai, China; 1:5,000) and *Nl*ANX5-YFP protein was detected by using an anti-GFP antibody (catalog no. M20004, Abmart, Shanghai, China; 1:5,000).

**RNA isolation and quantitative real-time PCR (qRT-PCR).** Total RNA was extracted using Trizol reagent (Invitrogen, Carlsbad, CA, USA) and RT-qPCR was performed as described previously (12). Briefly, 1.0 µL DNase-treated RNA was reverse-transcribed in a 20 µL reaction volume with oligo (dT) 18 primer using Moloney murine leukemia virus reverse transcriptase (Fermentas, Thermo Fisher Scientific, Waltham, MA, USA). Then, 1.0 µL cDNA was used for RT-qPCR using the SYBR Premix Ex Taq kit (Takara, Dalian, China) in a Roche LightCycler 480 real-time PCR machine, according to the manufacturer's instructions. At least three to four biological replicates for each genotype were used for RT-qPCR analysis. The *OsACTIN* gene (*LOC_Os03g50885*) was used as the control. All primers used for RT-qPCR are listed in Supplemental Table S2.

**AlphaFold3 (AF3) predictions.** The AF3-predicted models were generated using the AlphaFold Server (https://alphafoldserver.com/). The protein sequences for *Nl*ANX5 and *Os*ANN1–10 are listed in Supplementary Table S3. *Nl*ANX5 protein was paired with each *Os*ANN protein and analyzed under conditions with or without calcium ions. The number of calcium ions ranged from 0 to 12, and all predictions were consistently performed using a random seed value of 833366313 for comparison purposes.

**BPH honeydew measurement.** The total honeydew was collected by a pentahedron yurt-shaped Parafilm sachet (6 cm x 6 cm x 4 cm), which wrapped the stem of 6-leaf-stage rice seedlings. This Parafilm sachet was punched with around 30 breathable air holes on each of the four sides (excluding the bottom) using 1 ml syringes. Twenty ds*GFP*- and ds*NlANX5*-treated BPH nymphs in the end 4$^{th}$ instar were recovered for 24 h as described previously and the larvae in the early 5$^{th}$ instar were confined in the Parfilm sachet as a biological replicate. The experiment was conducted under the climate-controlled environment with 25 ± 0.5°C temperature, 50 ± 0.5% relative humidity and a photoperiod of 16 h of illumination and 8 h of darkness. The Parafilm sachets with or without honeydew were weighed before and after BPH fed for 24 h using an electronic balance (0.0001 g sensitivity, Meilen, Shenzhen, China). The experiment has at least 10 replicates for each treatment and the relative honeydew weights were calculated by the formula as follows:

$$H_{Rw} = \frac{W_{BHW}}{W_{BIW}}$$

$H_{RW}$: the relative honeydew weight of BPH after feeding for 24h; $W_{BHW}$: the honeydew weight of BPH after feeding for 24h; $W_{BIW}$: the initial weight of BPH before feeding.

**Phytohormone detection and quantification.** Rice samples were collected from the six-leaf stage rice seedlings (Nip and *OsANX5*-OE) before and after ds*GFP* and ds*NlANX5* BPH fed for 48 h, Mock were plants infested by BPH without injection. The three outermost layers of rice leaf sheath surrounding the stem were cut into small pieces (around 0.5 cm) and flash-frozen immediately. The frozen rice leaf tissues were ground into powder under liquid nitrogen condition and 100 mg powder was used for

LC-MS. Quantification of plant hormones/metabolites were performed at the Shanghai Hoogen-biology Limited Company of China using AB Sciex ExionLC Liquid Chromatography and AB6500plus Mass Spectrometer. The phytohormone standard solution was tested by LC-MS/MS to examine the linear range and generate the standard curve. The correlation coefficient of the linear regression equation was over 0.99. The external standard method was used to quantify the target sample concentration by the specific calculation formula as follows:

$$C_{SPL} = \frac{A_{SPL} - b}{k}$$

$C_{SPL}$: the target concentration in the sample solution; $A_{SPL}$: the peak area of the target ion in the sample solution; $k$: the slope in the standard curve; $b$: the intercept in the standard curve. The final sample quantitative concentration is calculated based on the standard curve and the target signal intensity detected in the sample.

### Figures S1 to S17

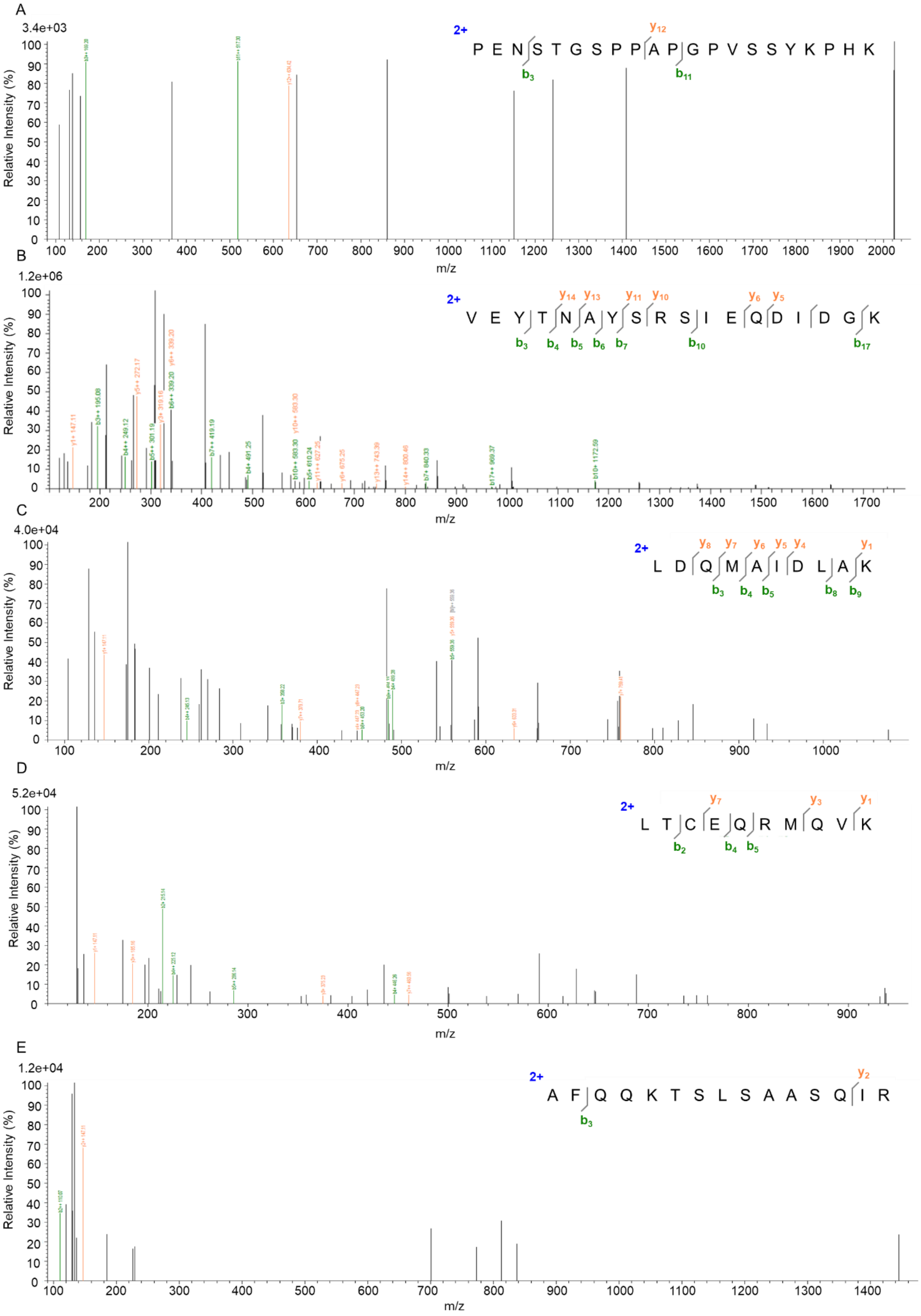

F

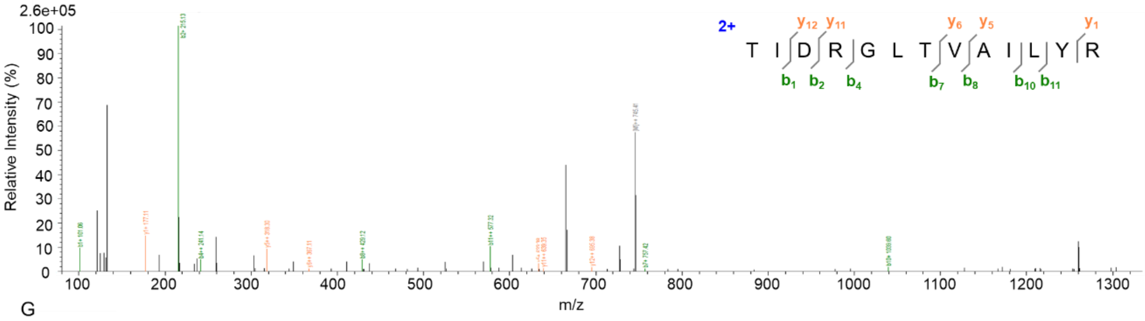


G

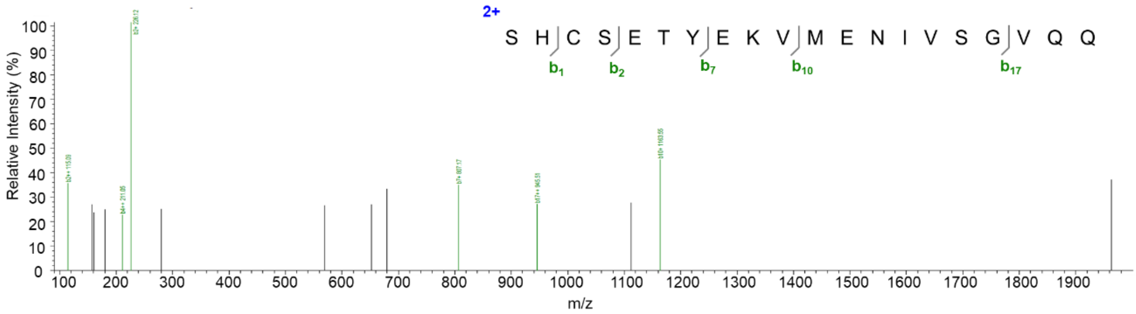


**Fig. S1. B & Y ion data of seven *Nl*ANX5 peptide fragments detected in leaf sheaths by LC-MS, related to Figure 1*B*.** (*A*–*G*) These seven figure panels show matches of the detected secondary ions to specific peptide sequence fragments of *Nl*ANX5. The Y axis represents relative intensity of ion (%), while X axis denotes mass-to-charge ratio distribution of detected secondary ions (m/z).

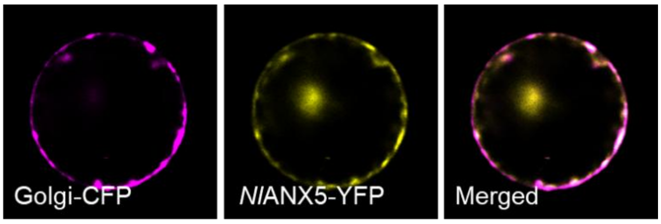


**Fig. S2. Subcellular localization of *Nl*ANX5, related to Figure 1C.** *Nl*ANX5 is colocalized with the YFP signals in the Golgi of rice cells. Golgi-CFP and *Nl*ANX5-YFP fusion protein were co-expressed in rice protoplasts 16 h after the corresponding DNA constructs were introduced into rice protoplasts via polyethylene glycol-mediated transformation. The Golgi marker was the cytoplasmic tail and transmembrane domain of soybean α-1,2-mannosidase 1 (*Gm*Man1).

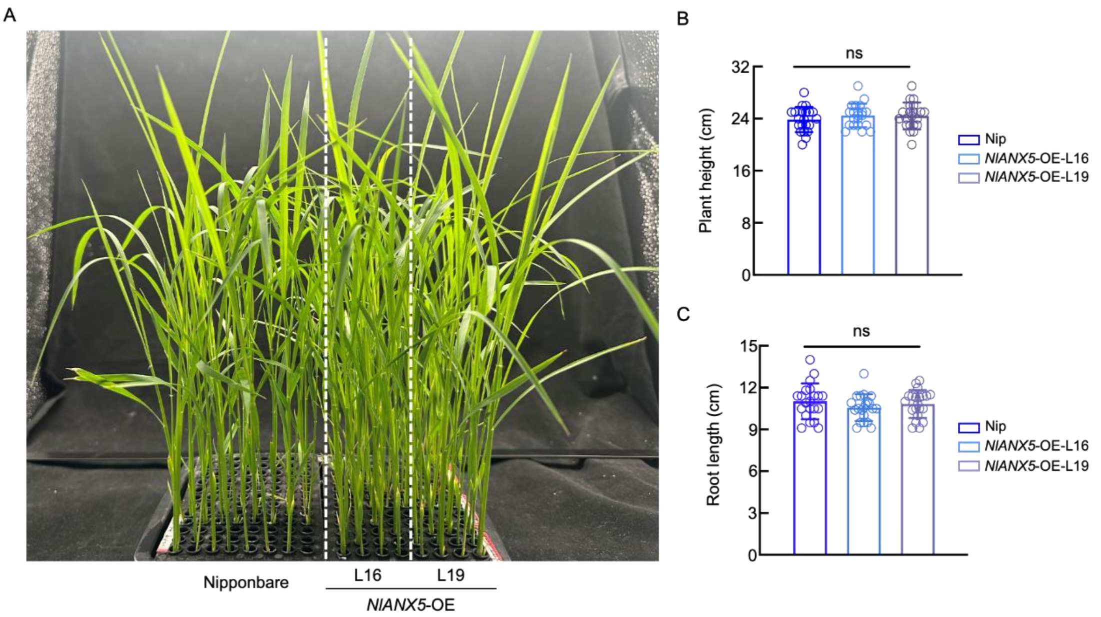


**Fig. S3. Growth phenotypes of *NlANX5*-expressing rice plants, related to Figure 2.** (*A*) 21-day-old of Nipponbare, *NlANX5*-OE L16 and *NlANX5*-OE L19 plants grown in the modified Yoshida medium. *NlANX5*-OE L16 and *NlANX5*-OE L19 plants displayed normal appearance compared with Nipponbare. Photos were taken after 4-week growth of seedlings. (*B* and *C*) Plant height (*B*) and root length (*C*), respectively, in (*A*). Values are displayed as mean ± SD (*n* = 20 plants). ns indicate no significant statistically differences analyzed by one-way ANOVA (Tukey test).

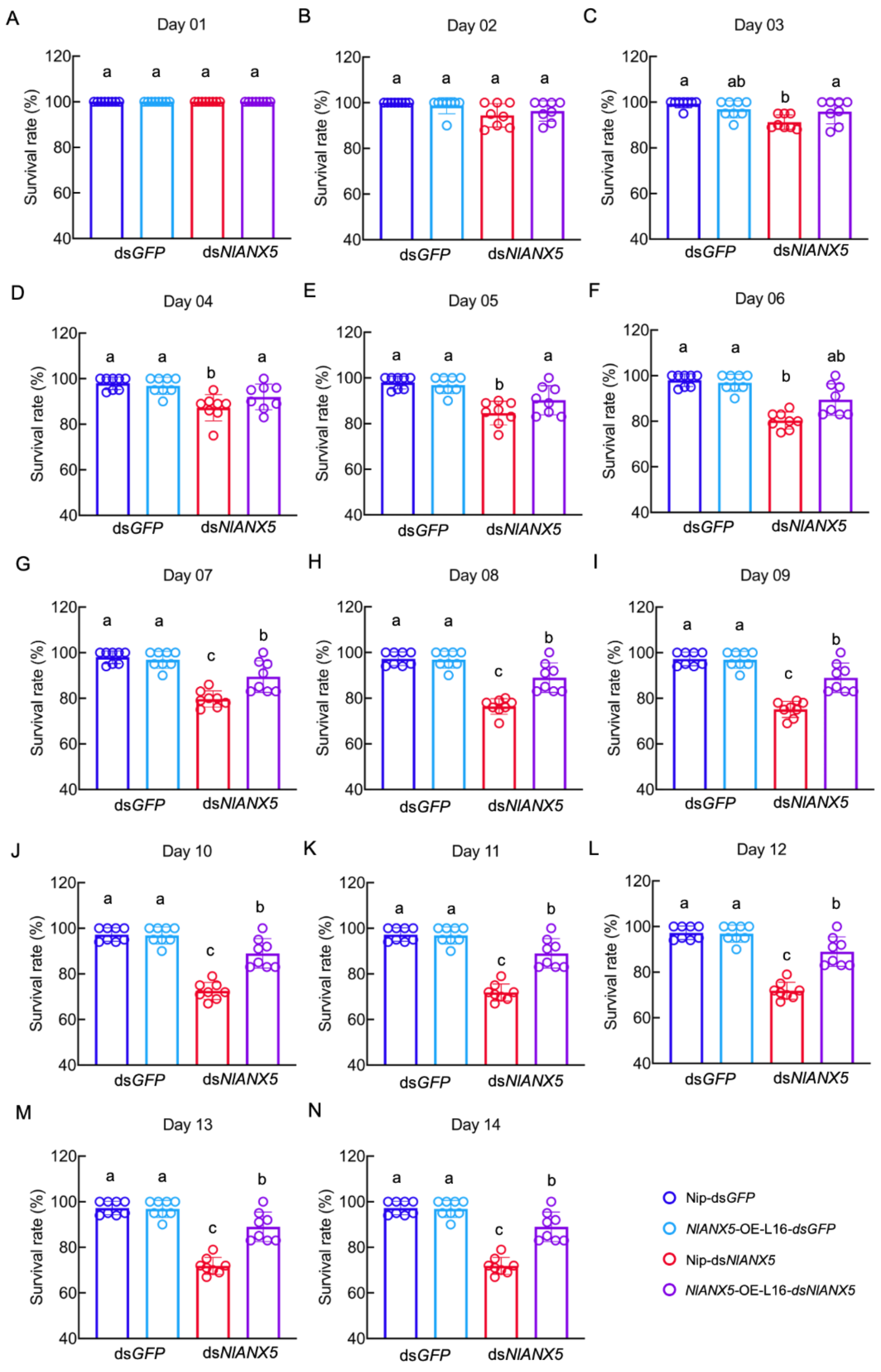


**Fig. S4. The daily survival rates of ds*GFP* and ds*NlANX5* BPH insects on Nipponbare and *NlANX5* transgenic plants line16, related to Figure 2.** (*A*–*N*) Values are represented as mean ± SEM (*n* > 6 biological replicates; 20 individual insects per each biological replicate). Different letters indicate statistically significant differences analyzed by two-way ANOVA (Tukey test, $P < 0.05$). Experiments were repeated three times.

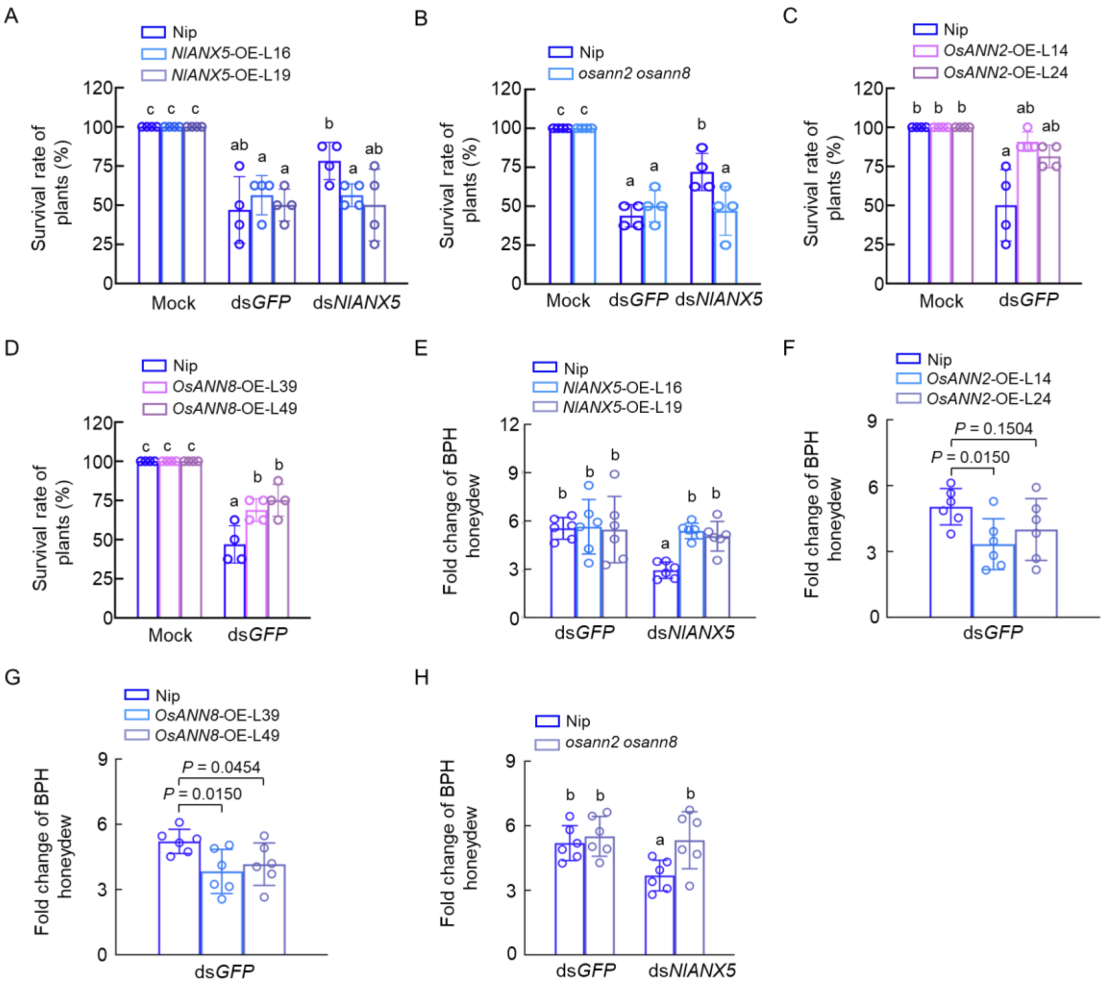


**Fig. S5. Survival rate of rice plants and BPH honeydew production related to Figure 2, 5 and 6** (*A*–*B*) Survival rate of 3-week-old Nip, *NlANX5*-expressing and *osann2 osann8* rice plants infested by 30 5$^{th}$ instar ds*GFP* and ds*NlANX5* BPH nymphs for 7 days. (*C*–*D*) Survival rate of 3-week-old Nip, *OsANN2*-expressing and *OsANN8*-expressing rice plants infested by 30 5$^{th}$ instar ds*GFP* and ds*NlANX5* BPH for 7 days. The mock treatment rice plants without BPH infestation were used as the control group. Values are displayed as mean ± s.d. (n = 4 replicates, each replicate including eight rice plants) and different letters indicate statistically significant differences analyzed by two-way ANOVA (Tukey test, $P < 0.05$). (*E*) Fold change of BPH honeydew over the initial weight of ds*GFP* and ds*NlANX5* BPH that feed on Nip and *NlANX5*-expressing rice plants. (*F*–*G*) Fold change of BPH honeydew over the initial weight of ds*GFP* BPH that feed on Nip, *OsANN2*-expressing rice plants and *OsANN8*-expressing rice plants. (*H*) Fold change of BPH honeydew of ds*GFP* BPH and ds*NlANX5* BPH on Nip and *osann2 osann8* rice plants. Values are displayed as mean ± s.d. (n = 20 biological replicates per treats, each treat has six repeats). Different letters indicate statistically significant differences analyzed by two-way ANOVA ($P < 0.05$), and *P* value above bars in *F*–*G* were calculated by student's *t* test.

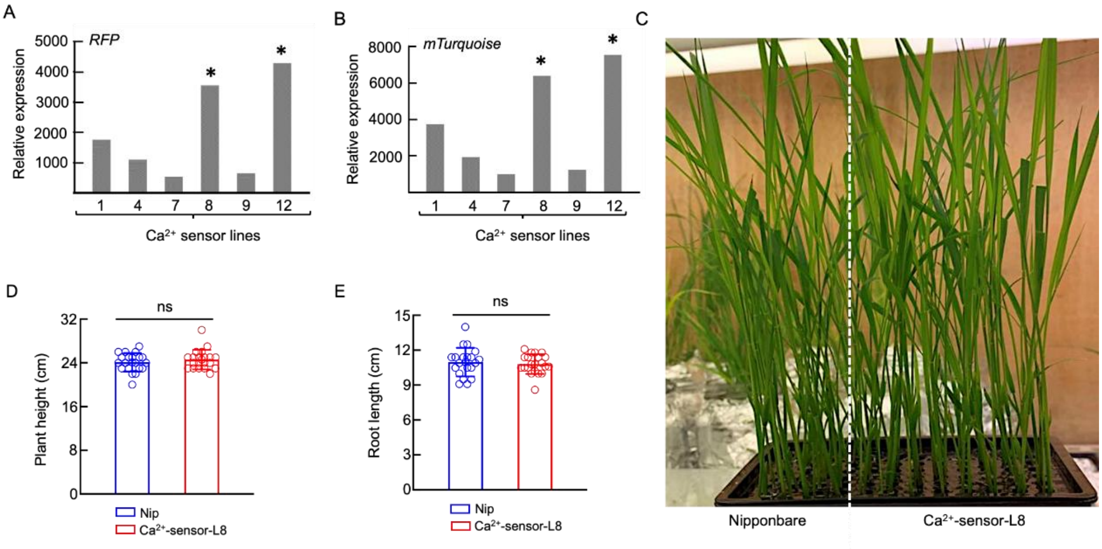


**Fig. S6. Screening of transgenic plants expressing a ratiometric cytoplasmic $Ca^{2+}$-sensor in background of Nipponbare, related to Figures 3*A* and 3*B*.** (*A*–*B*) Transcript levels of *RFP* (*A*) and *mTurquoise* (*B*) in the cyto-$Ca^{2+}$-sensor expressing rice plants. Six independent transgenic rice lines were selected to determine the relative expression of *RFP* and *mTurquoise*, the line labeled with the asterisk was used for further calcium level assays. (*C*) Growth phenotypes of Nipponbare and cyto-$Ca^{2+}$-sensor expressing rice plants (T3 generation). Photos were taken after 4-week growth of seedlings. (*D* and *E*) Plant height (*D*) and root length (*E*), respectively, in (*C*). Values are displayed as mean ± SD (*n* = 20 plants). ns indicate no significant statistically differences analyzed by one-way ANOVA (Tukey test).

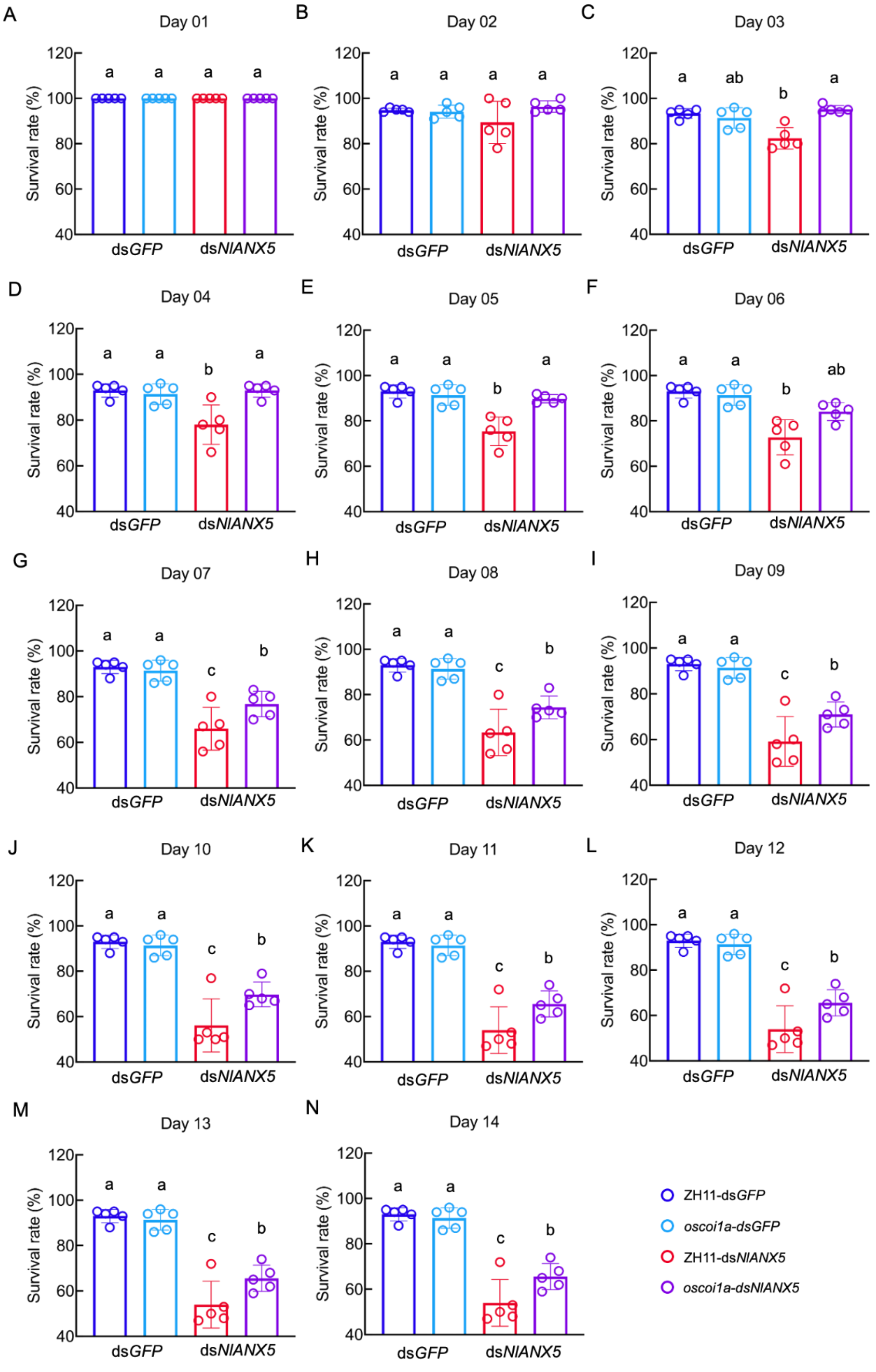


**Fig, S7. The daily survival rates of ds*GFP* and ds*NlANX5* injected BPH insects feeding on ZH11 or *oscoi1a* mutant plants, related to Figure 3.** Values are displayed as mean ± s.e.m. (*n* = 5 biological replicates for each genotype), and different letters indicate statistically significant differences analyzed by two-way ANOVA (Tukey test, $P < 0.05$). The 2$^{nd}$ instar nymphs were used in the survival assays. Results were repeated three times with similar trends.

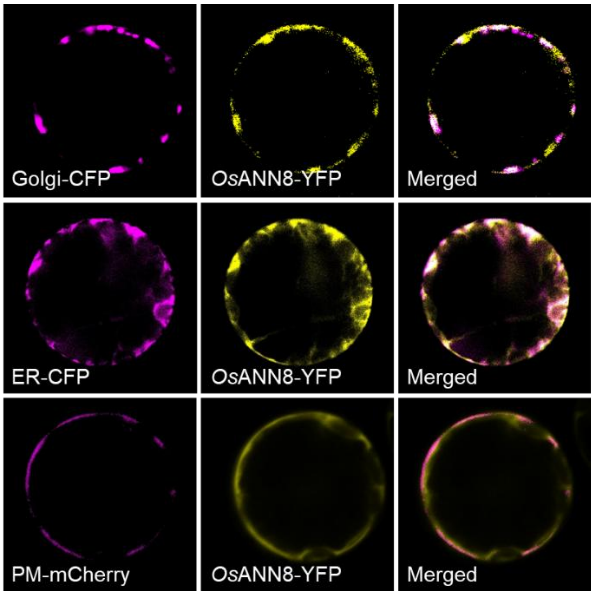


**Fig. S8. Subcellular localization of *Os*ANN8 in rice protoplasts, related to Figure 4.** *Os*ANN8 is colocalized with the YFP signals in the Golgi (upper row), ER (middle row) and plasm membrane (PM) (lower row) of rice cells. Golgi-CFP and *Os*ANN8-YFP, ER-CFP and *Os*ANN8-YFP, or PM-mCherry and *Os*ANN8-YFP fusion proteins were detected in rice protoplasts 16 h after the corresponding DNA constructs were introduced into rice protoplasts via polyethylene glycol-mediated transformation. Experiments were repeated three times.

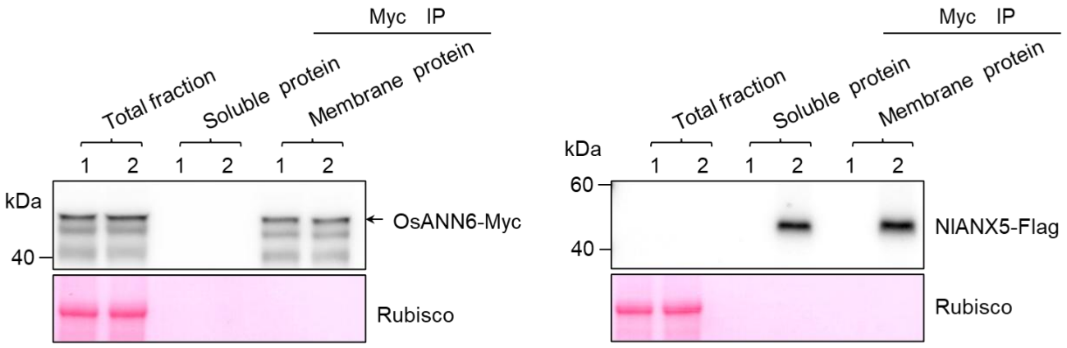


**Fig. S9 Effects of *Nl*ANX5 on the association of *Os*ANN6 with the rice membrane.** Sample 1 represents rice protoplasts expressing *Os*ANN6-Myc alone, whereas sample 2 contains rice protoplasts co-expressing *Os*ANN6-Myc with *Nl*ANX5-Flag. Soluble and membrane proteins were immunoprecipitated by both Myc-trap and Flag-trap beads, respectively, to detect *Os*ANN6-Myc and *Nl*ANX5-Flag proteins. Experiments were repeated three times with similar trends.

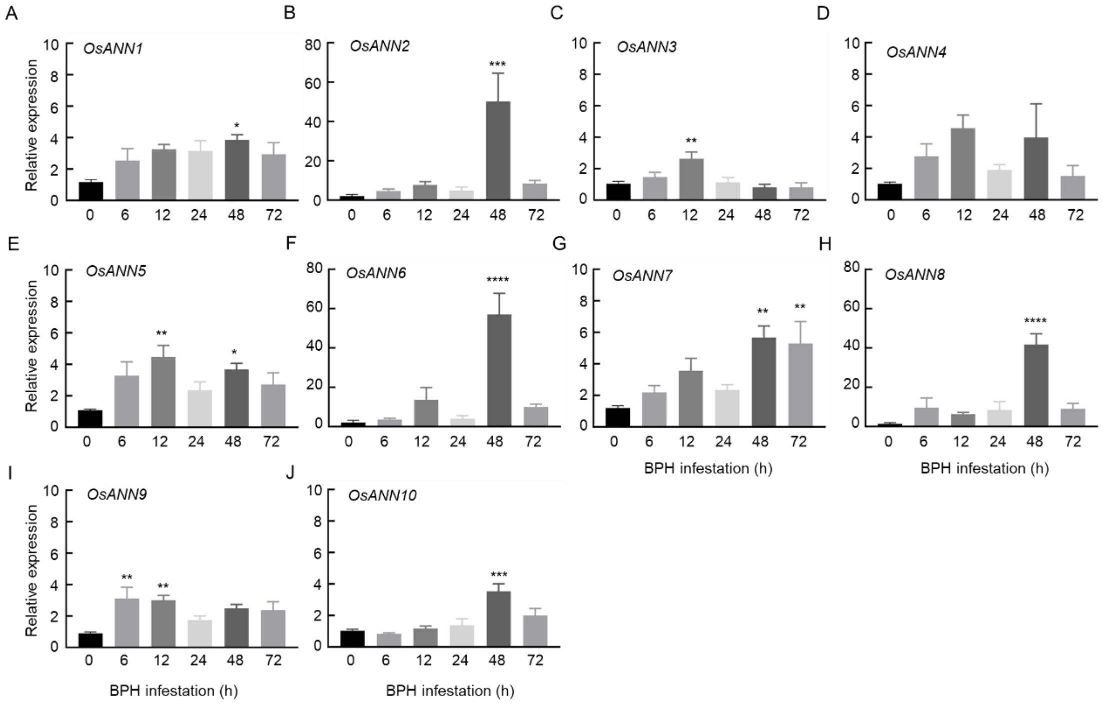


**Fig. S10. Expression pattern of *OsANN* genes in Nipponbare plants before and after BPH feeding, related to Figure 5.** Four-leaf-stage Nipponbare plants were fed by BPH and RNA samples were collected in the indicated time. Values are displayed as mean ± s.e.m. ($n$ = 3 biological replicates). The asterisk(s) represent statistically significant differences between the indicated infestation time with 0 h analyzed by student's $t$-test (*$P$ < 0.05, **$P$ < 0.01, ***$P$ < 0.001, ****$P$ < 0.0001). The *OsACTIN* (*Os03g50885*) gene was used as internal control.

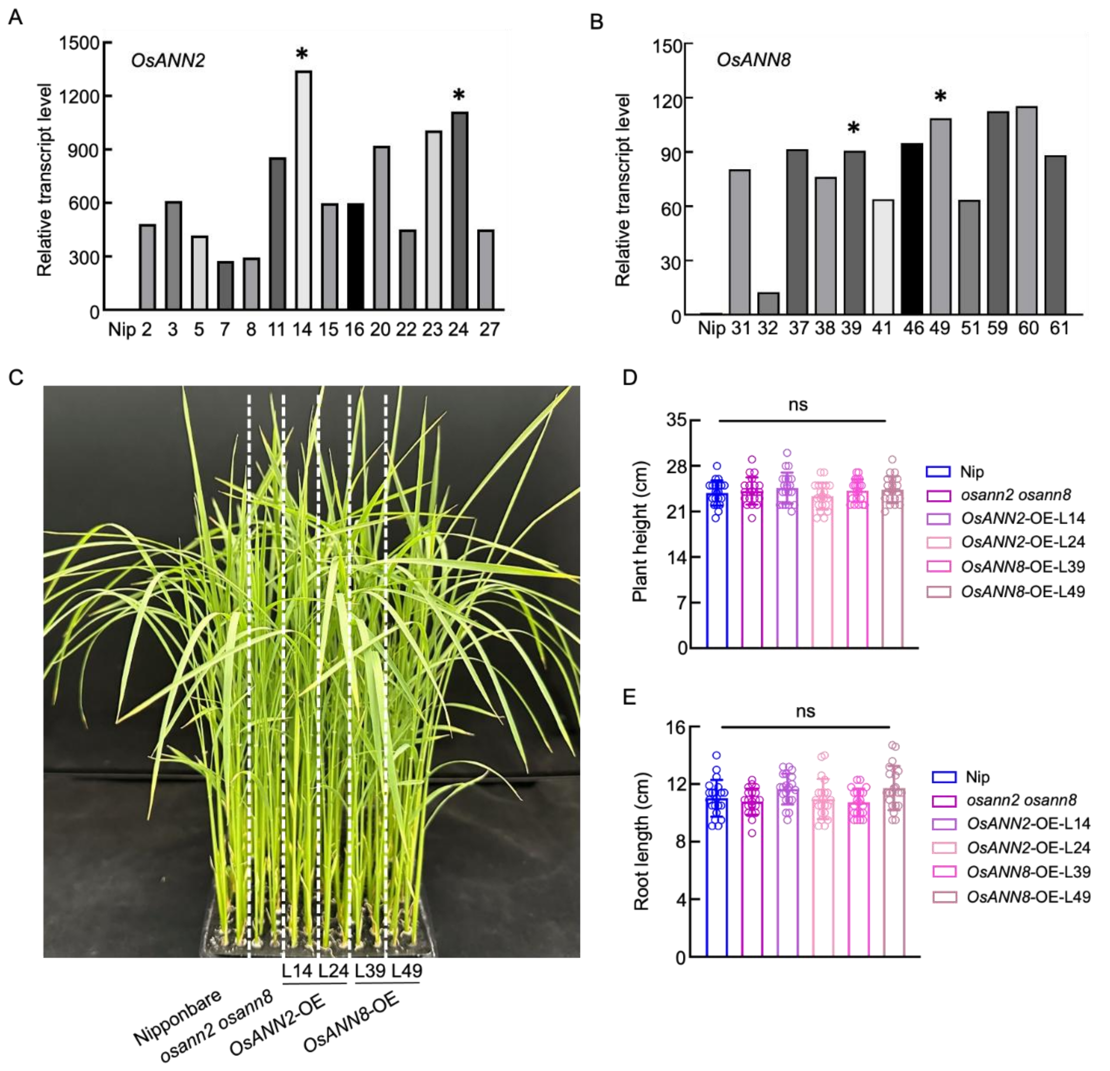


**Fig. S11. Screening of *OsANN2*-OE and *OsANN8*-OE transgenic rice plants, related to Figure 5.** (*A*) Transcript level of *OsANN2* in the *OsANN2-OE* transgenic plants. The lines labeled with the asterisk were used for further assays. (*B*) Transcript level of *OsANN8* in the *OsANN8-OE* transgenic plants. The lines labeled with the asterisk were used for further assays. RNA samples were collected from 3-week-old T3 transgenic plants. Values are displayed as mean ± s.e.m. ($n$ = 3 biological replicates). The *OsACTIN* (*Os03g50885*) gene was used as internal control. (*C*) Growth phenotypes of *OsANN2*- and *OsANN8*-expressing rice plants compared with control Nipponbare plants. Photos were taken after 3-week growth. (*D* and *E*) Plant height (*D*) and root length (*E*), respectively, in (*C*). Values are displayed as mean ± SD ($n$ = 20 plants). ns indicate no significant statistically differences analyzed by one-way ANOVA (Tukey test).

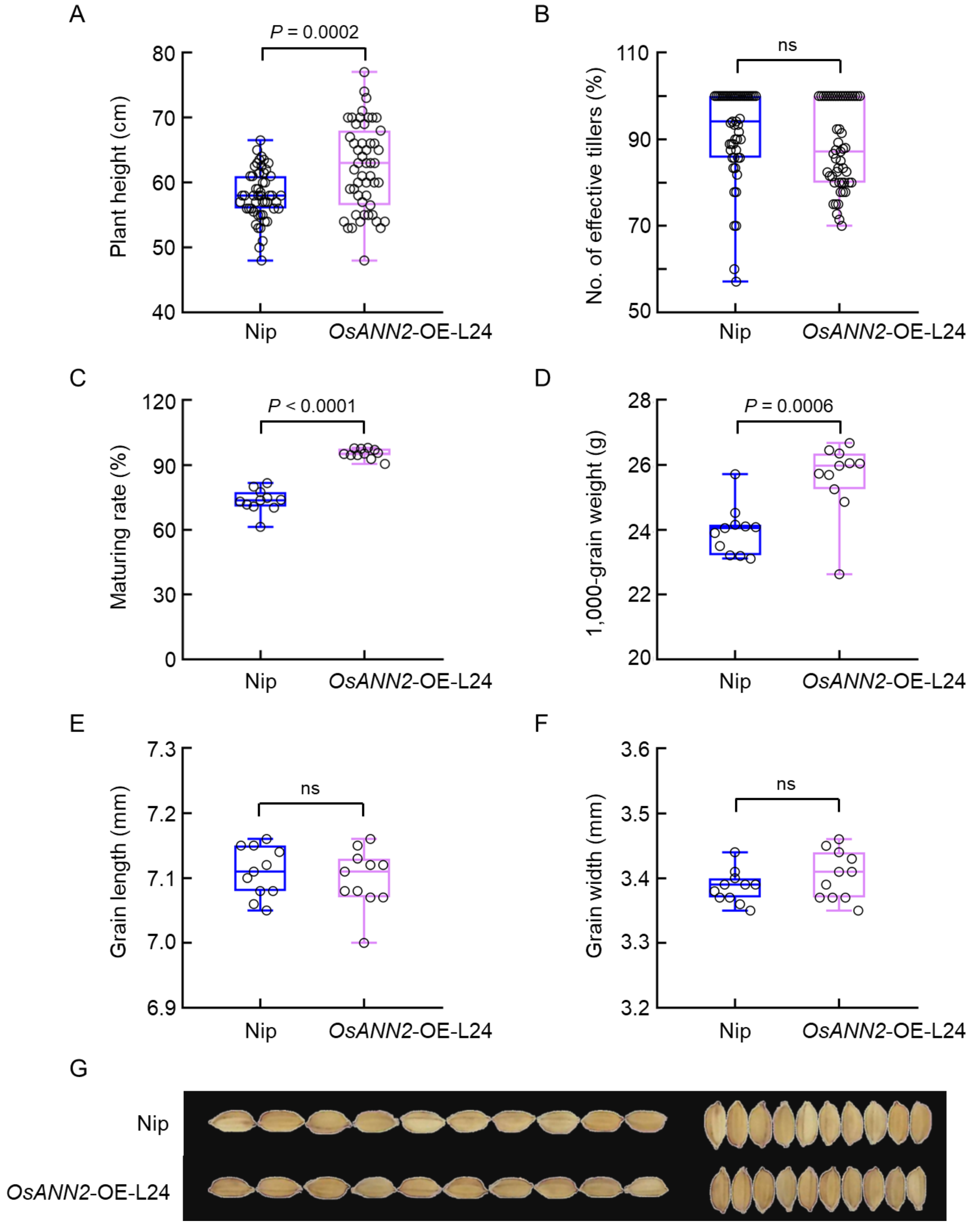


**Fig. S12. Agronomic traits of *OsANN2*-expressing rice plants, related to Figure 5.** Plant height (*A*), percentage of effective tillers (*B*), percentage of maturing rate (*C*), 1,000-grain weight (*D*), grain length (*E* and *G*) and grain weight (*F* and *G*) of Nip and *OsANN2*-OE-L24. Values are displayed as mean ± s.d., ns indicates no statistically significant differences analyzed by unpaired student *t*-test.

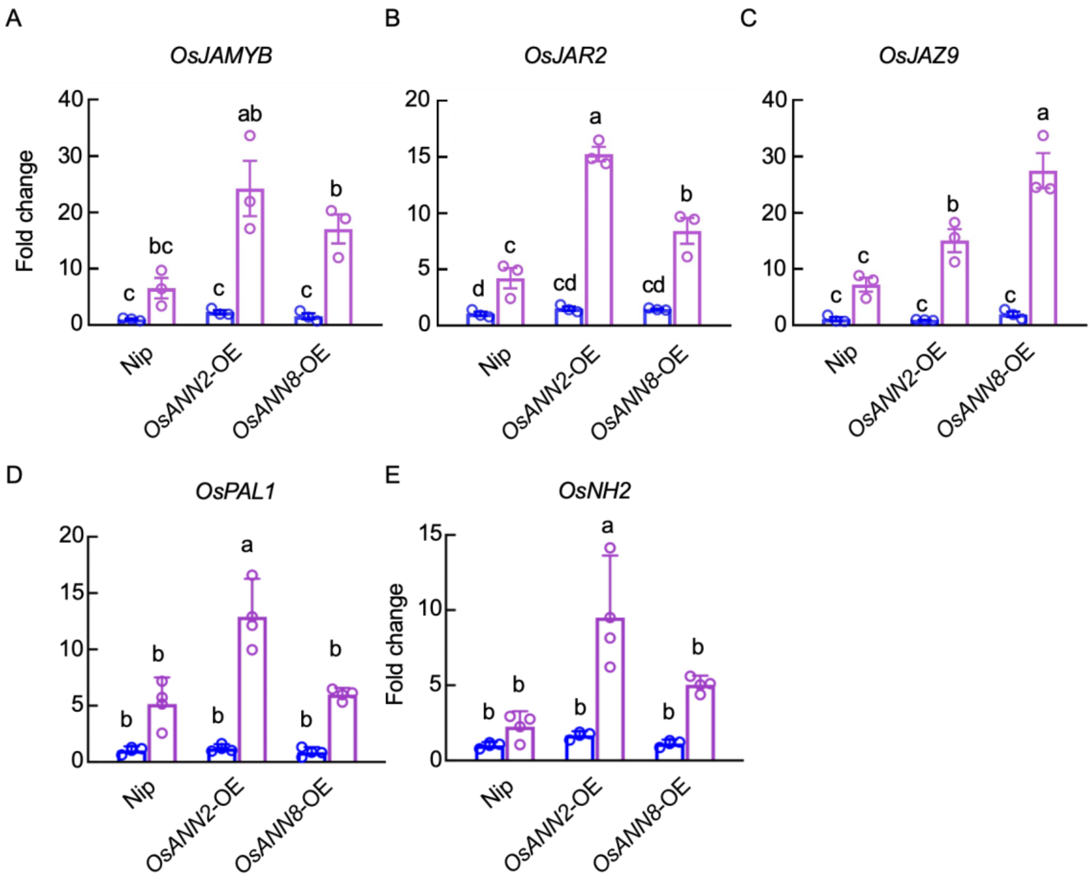


**Fig. S13. Transcript levels of JA and SA biosynthesis as well as responsive genes, related to Figure 5.** (*A–C*) Transcript levels of rice JA biosynthesis and responsive genes, such as *OsJAMYB*, *OsJAR2* and *OsJAZ9* in Nip, *OsANN2-OE* and *OsANN8-OE* plants. (*D–E*) Transcript levels of rice SA biosynthesis gene, *OsPAL1* and *OsNH2* in Nip and *OsANN2-OE* and *OsANN8-OE* plants. The *OsACTIN* (*Os03g50885*) gene was used as internal control. Each 5-leaf-period rice plant was treated with 20 fifth instar BPH for 24 h. Values are displayed as mean ± s.e.m. ($n \geq 3$ biological replicates). Different letters indicate statistically significant differences analyzed by two-way ANOVA (Tukey test, $P < 0.05$). Experiments were repeated three times.

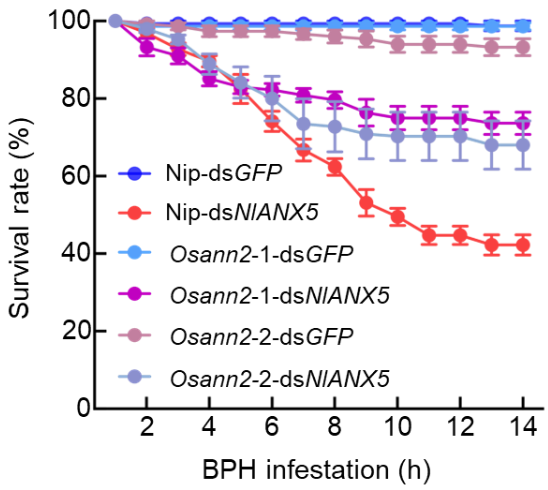


**Fig. S14. Partial restoration of the survival defect of ds*NlANX5* BPH in *Osann2* single mutant plants, related to Figure 6.** The survival rates of ds*GFP* and ds*NlANX5* BPH insects fed on Nip and *osann2* single mutant plants. Values are represented as mean ± s.e.m. of 8 biological replicates (Two-way ANOVA; 20 individual 2$^{nd}$ BPH nymphs per rice plant for each biological replicate).

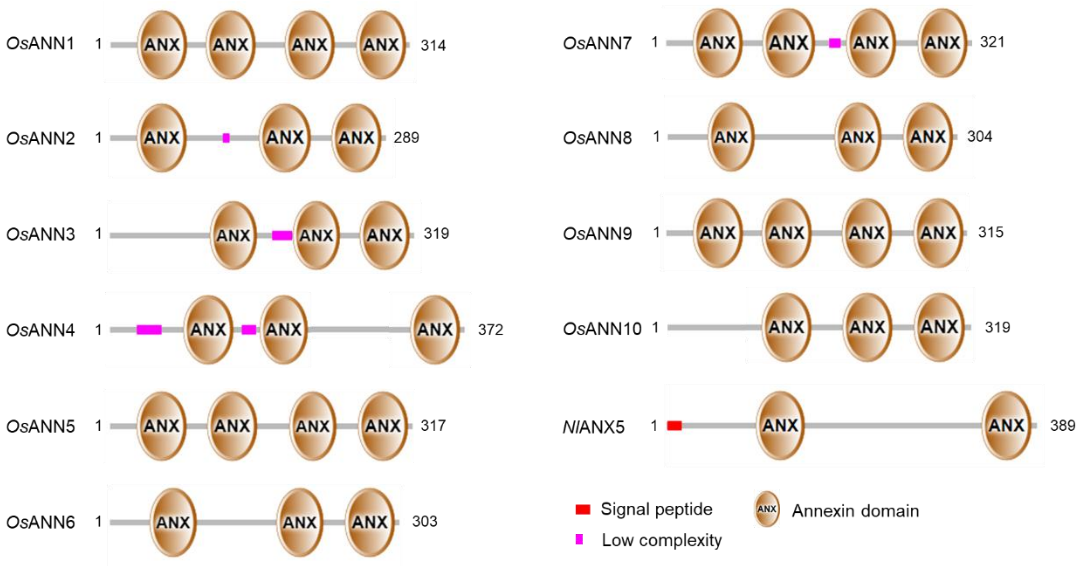


**Fig. S15. The predicted domain structures of *Nl*ANN5 and *Os*ANNs by SMART (Simple Modular Architecture Research Tool).** Red rectangle indicates the putative signal peptide. Megata rectangle means low complexity area. Brown ovals represent the annexin domain(s). The size bar shows the amino acid residues of the deduced proteins. *Nl*ANX5: ANJ04663.1, *Os*ANN1: LOC_Os02g51750.1, *Os*ANN2: LOC_Os01g31270.1, *Os*ANN3: LOC_Os05g31750.1, *Os*ANN4: LOC_Os05g31760.1, *Os*ANN5: LOC_Os06g11800.1, *Os*ANN6: LOC_Os07g46550.1, *Os*ANN7: LOC_Os08g32970.1, *Os*ANN8: LOC_Os09g20330.1, *Os*ANN9: LOC_Os09g23160.1 and *Os*ANN10: LOC_Os09g27990.1.

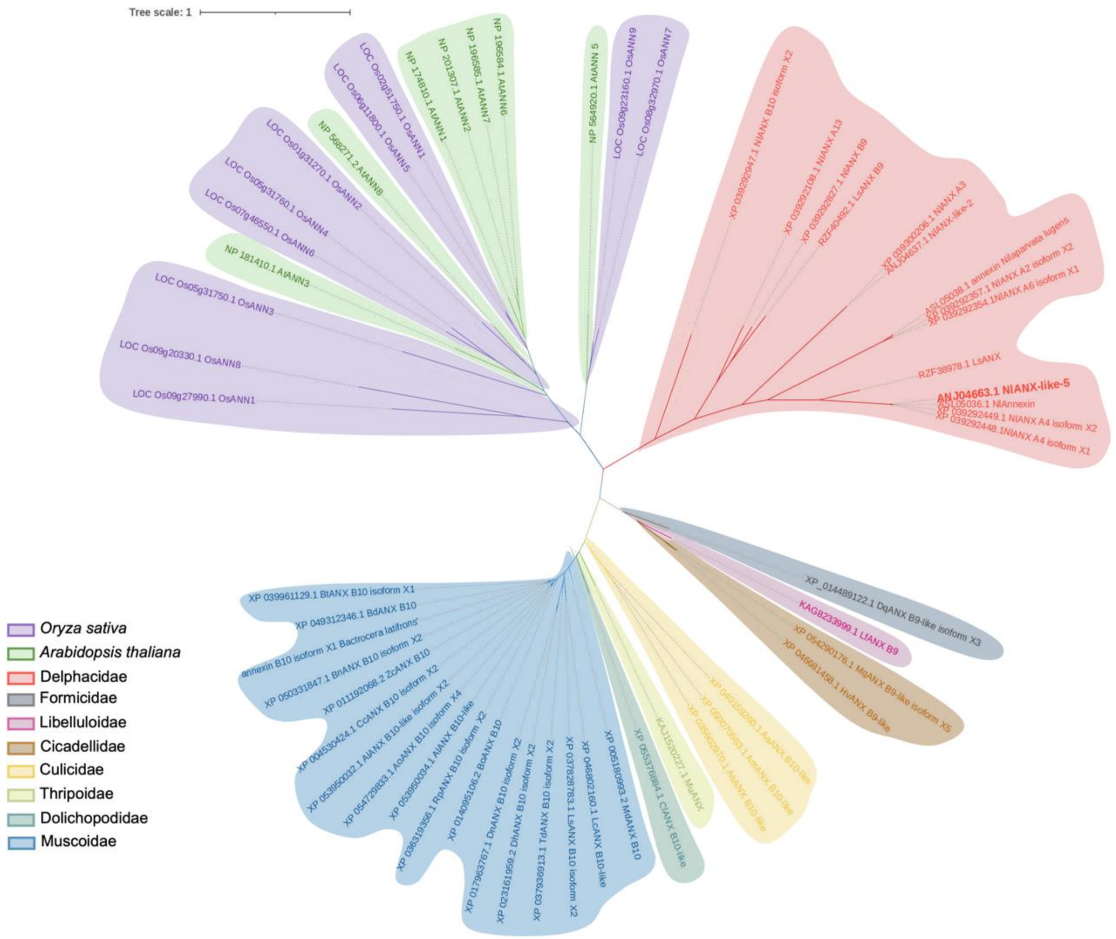


**Fig. S16. Phylogenic analysis of annexin proteins from representative insect and plant species.** The protein sequences were obtained from NCBI (www.ncbi.nlm.nih.gov), TAIR (http://www.arabidopsis.org/) and Rice Genome Annotation Project (http://rice.uga.edu/index.shtml). The bold red branch denotes as *Nl*ANX5 (AN J04663.1 *Nl*ANX-like-5). Sequence alignment was performed using Clustal W, and the phylogenetic tree was generated by MEGA 11.0 using the Maximum-likelihood method. Bootstrap values from 1,000 replicates were used to assess the robustness of the tree. The phylogenic tree was beautified with iTOL V6 (https://itol.embl.de/). The scale indicates the average number of substitutions per site.

Figure 1D

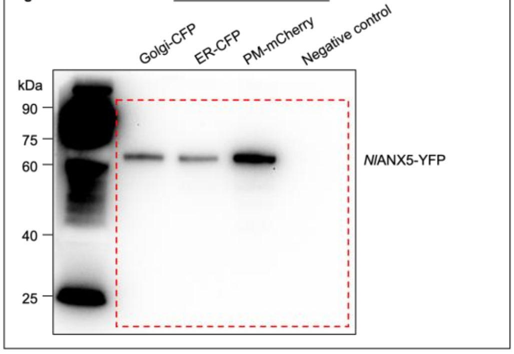


Figure 4A

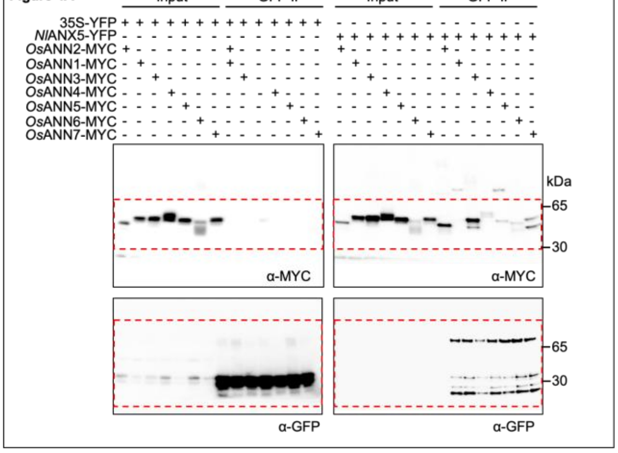


Figure 4B

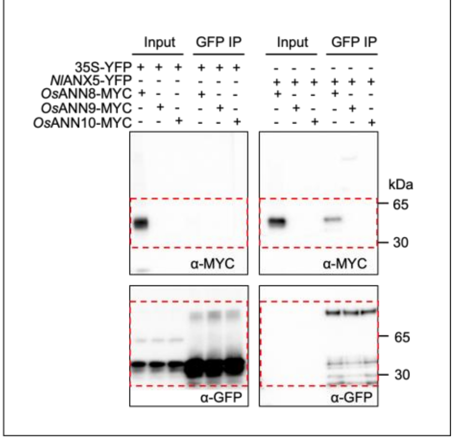


Figure 4E

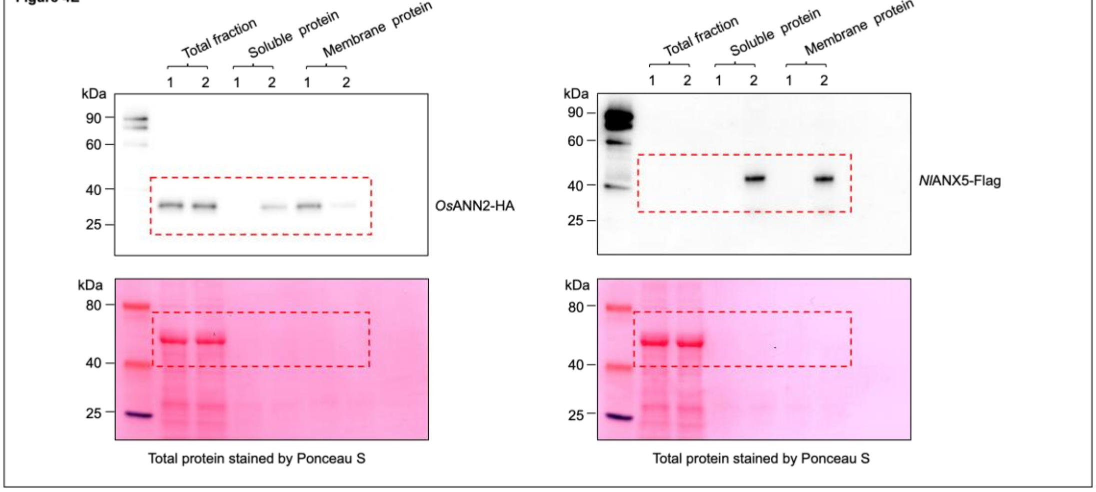

**Continued.**

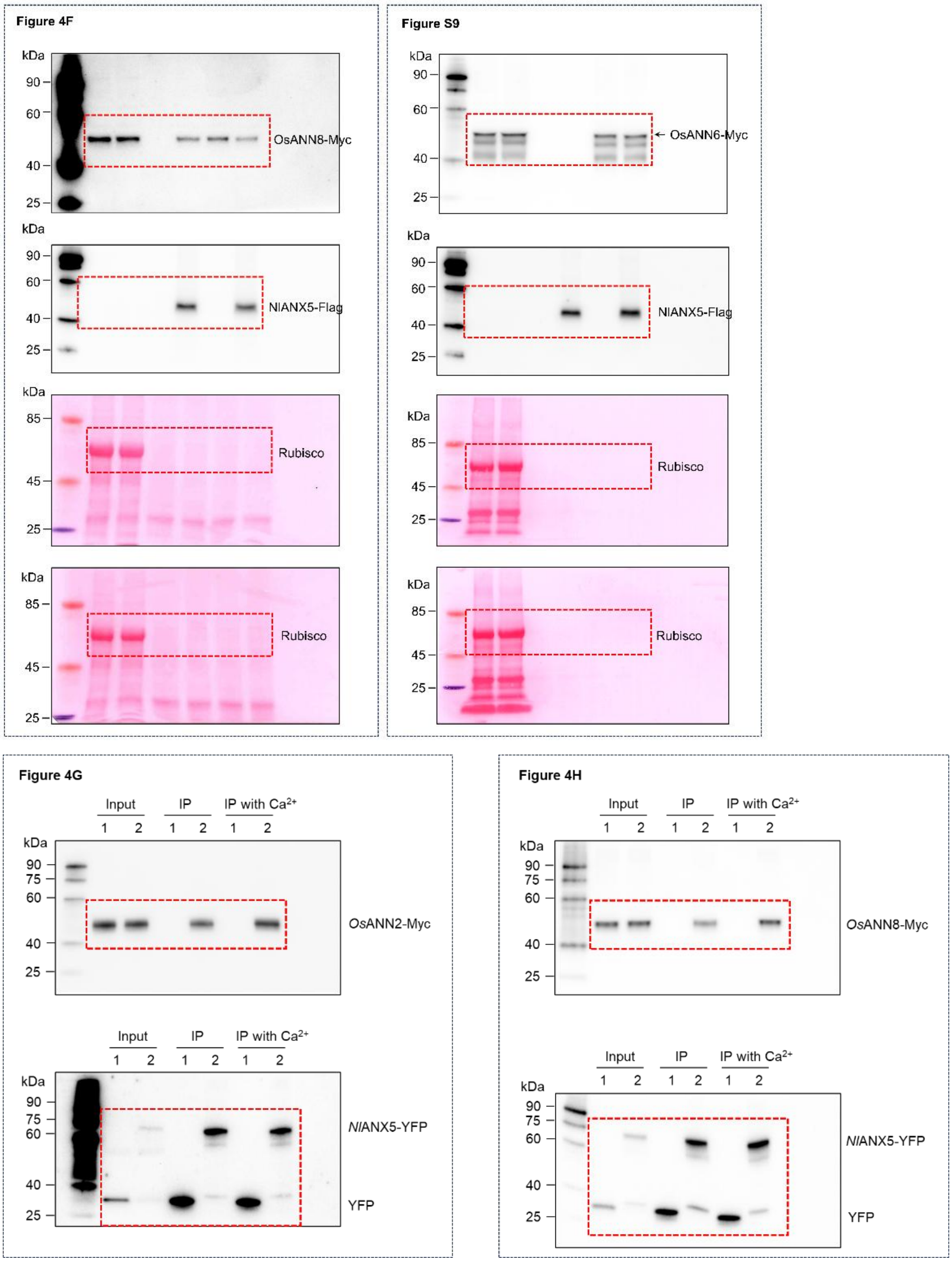


**Fig. S17. Uncropped images for protein gels, related to Figure 1*D*, Figure 4*A–B* and Figure 4*E–H* and Figure S9.**

**Table S1. *Nl*ANX5-*Os*ANN interactions predicted by AlphaFold3 in the absence or presence of calcium ions, related to Figure 4.**

| $Ca^{2+}$ ions | 0 | 1 | 2 | 3 | 4 | 5 | 6 | 7 | 8 | 9 | 10 | 11 | 12 |
|---|---|---|---|---|---|---|---|---|---|---|---|---|---|
| **ipTM** | | | | | | | | | | | | | |
| *Os*ANN1 | 0.22 | 0.27 | 0.22 | 0.22 | 0.23 | 0.20 | 0.18 | 0.19 | 0.19 | 0.16 | 0.19 | 0.16 | 0.13 |
| ***Os*ANN2** | **0.19** | **0.45** | **0.46** | **0.44** | **0.22** | **0.23** | **0.47** | **0.32** | **0.57** | **0.29** | **0.24** | **0.55** | **0.24** |
| *Os*ANN3 | 0.11 | 0.12 | 0.12 | 0.12 | 0.13 | 0.12 | 0.13 | 0.13 | 0.14 | 0.15 | 0.13 | 0.14 | 0.14 |
| *Os*ANN4 | 0.16 | 0.17 | 0.19 | 0.18 | 0.16 | 0.17 | 0.23 | 0.16 | 0.16 | 0.16 | 0.16 | 0.16 | 0.16 |
| *Os*ANN5 | 0.15 | 0.20 | 0.22 | 0.21 | 0.21 | 0.19 | 0.18 | 0.19 | 0.25 | 0.26 | 0.26 | 0.29 | 0.27 |
| ***Os*ANN6** | **0.14** | **0.17** | **0.21** | **0.24** | **0.22** | **0.58** | **0.21** | **0.18** | **0.20** | **0.20** | **0.36** | **0.33** | **0.43** |
| *Os*ANN7 | 0.44 | 0.28 | 0.21 | 0.25 | 0.22 | 0.17 | 0.23 | 0.14 | 0.15 | 0.12 | 0.22 | 0.19 | 0.20 |
| ***Os*ANN8** | **0.37** | **0.32** | **0.30** | **0.31** | **0.37** | **0.28** | **0.33** | **0.34** | **0.28** | **0.31** | **0.71** | **0.28** | **0.71** |
| *Os*ANN9 | 0.26 | 0.33 | 0.20 | 0.19 | 0.29 | 0.50 | 0.42 | 0.49 | 0.36 | 0.36 | 0.31 | 0.30 | 0.29 |
| *Os*ANN10 | 0.15 | 0.27 | 0.20 | 0.16 | 0.15 | 0.20 | 0.17 | 0.19 | 0.16 | 0.17 | 0.17 | 0.16 | 0.17 |
| **pTM** | | | | | | | | | | | | | |
| *Os*ANN1 | 0.55 | 0.57 | 0.55 | 0.55 | 0.55 | 0.54 | 0.53 | 0.53 | 0.53 | 0.52 | 0.53 | 0.52 | 0.51 |
| ***Os*ANN2** | **0.55** | **0.61** | **0.61** | **0.61** | **0.56** | **0.57** | **0.61** | **0.57** | **0.69** | **0.56** | **0.56** | **0.66** | **0.55** |
| *Os*ANN3 | 0.50 | 0.51 | 0.51 | 0.51 | 0.51 | 0.51 | 0.51 | 0.51 | 0.51 | 0.52 | 0.51 | 0.51 | 0.51 |
| *Os*ANN4 | 0.50 | 0.50 | 0.51 | 0.51 | 0.50 | 0.50 | 0.52 | 0.50 | 0.50 | 0.50 | 0.50 | 0.50 | 0.50 |
| *Os*ANN5 | 0.52 | 0.54 | 0.55 | 0.54 | 0.54 | 0.54 | 0.53 | 0.53 | 0.56 | 0.56 | 0.56 | 0.57 | 0.57 |
| ***Os*ANN6** | **0.52** | **0.53** | **0.55** | **0.56** | **0.55** | **0.70** | **0.55** | **0.53** | **0.54** | **0.54** | **0.61** | **0.59** | **0.64** |
| *Os*ANN7 | 0.64 | 0.57 | 0.54 | 0.56 | 0.55 | 0.53 | 0.53 | 0.51 | 0.52 | 0.50 | 0.54 | 0.53 | 0.53 |
| ***Os*ANN8** | **0.56** | **0.56** | **0.55** | **0.56** | **0.57** | **0.55** | **0.56** | **0.57** | **0.55** | **0.56** | **0.70** | **0.55** | **0.68** |
| *Os*ANN9 | 0.55 | 0.59 | 0.53 | 0.53 | 0.56 | 0.68 | 0.63 | 0.68 | 0.59 | 0.59 | 0.57 | 0.57 | 0.56 |
| *Os*ANN10 | 0.52 | 0.58 | 0.54 | 0.53 | 0.52 | 0.55 | 0.54 | 0.54 | 0.53 | 0.53 | 0.53 | 0.53 | 0.53 |

Summary of the interface predicted template modeling (ipTM; above) and the predicted template modeling (pTM; below) scores from AlphaFold3 analyses of the *Nl*ANX5-*Os*ANNs interactions with increasing number of calcium ions. The highest scores for *Os*ANN2, *Os*ANN6 and *Os*ANN8 are highlighted in yellow (Model seed: 833366313).

**Table S2. Primer sequences used in this study.**

| Primer name | Primer sequence (5’->3’) | Comments |
|---|---|---|
| *NlANX5*-pDONR207-F | ggggacaagtttgtacaaaaaagcaggcttcATGCT AGATGTTTCAGTATGTGTAT | Used for amplification of WT *NlANX5* |
| *NlANX5*-pDONR207-R | ggggaccactttgtacaagaaagctgggtcCTGCT GTACTCCACTGACG | Used for amplification of WT *NlANX5* |
| *OsANX1*-pDONR207-F | ggggacaagtttgtacaaaaaagcaggcttcATGG CGACGCTCACCGTG | Used for amplification of WT *OsANX1* |
| *OsANX1*-pDONR207-R | ggggaccactttgtacaagaaagctgggtcCTCTG CTCCAAGGAGGGC | Used for amplification of WT *OsANX1* |
| *OsANX2*-pDONR207-F | ggggacaagtttgtacaaaaaagcaggcttcATGG CAACGATAGTAGTTCCT | Used for amplification of WT *OsANX2* |
| *OsANX2*-pDONR207-R | ggggaccactttgtacaagaaagctgggtcGATGC CACTCCCAAGTAGAG | Used for amplification of WT *OsANX2* |
| *OsANX3*-pDONR207-F | ggggacaagtttgtacaaaaaagcaggcttcATGG CTGATGAAATCCAGCAT | Used for amplification of WT *OsANX3* |
| *OsANX3*-pDONR207-R | ggggaccactttgtacaagaaagctgggtcCTTGC CGCCGGCGACGAG | Used for amplification of WT *OsANX3* |
| *OsANX4*-pDONR207-F | ggggacaagtttgtacaaaaaagcaggcttcATGTG TTGCTGGTGCTGCTG | Used for amplification of WT *OsANX4* |
| *OsANX4*-pDONR207-R | ggggaccactttgtacaagaaagctgggtcCTTCTC AGGGCCGACGAG | Used for amplification of WT *OsANX4* |
| *OsANX5*-pDONR207-F | ggggacaagtttgtacaaaaaagcaggcttcATGG CGACGCTCACCGTC | Used for amplification of WT *OsANX5* |
| *OsANX5*-pDONR207-R | ggggaccactttgtacaagaaagctgggtcCTGCT CCTGACCCAGAAGA | Used for amplification of WT *OsANX5* |

| Primer name | Primer sequence (5’->3’) | Comments |
|---|---|---|
| *OsANX6-pDONR207-F* | ggggaccactttgtacaagaaagctgggtcCTGCT GTACTCCACTGACG | Used for amplification of WT *NlANX6* |
| *OsANX6*-pDONR207-R | ggggaccactttgtacaagaaagctgggtcGTCCT CCCTCCCGACCAA | Used for amplification of WT *NlANX6* |
| *OsANX7-pDONR207-F* | ggggaccactttgtacaagaaagctgggtcGCGGT CGCGGCCGATGAG | Used for amplification of WT *OsANX7* |
| *OsANX7-pDONR207-R* | ggggaccactttgtacaagaaagctgggtcCTCTG CTCCAAGGAGGGC | Used for amplification of WT *OsANX7* |
| *OsANX8-pDONR207-F* | ggggacaagtttgtacaaaaaagcaggcttcATGG CCTCGATTAGAGATTTTG | Used for amplification of WT *OsANX8* |
| *OsANX8-pDONR207-R* | ggggaccactttgtacaagaaagctgggtcATCTCT TGTCCTCAGGCCA | Used for amplification of WT *OsANX8* |
| *OsANX9*-pDONR207-F | ggggacaagtttgtacaaaaaagcaggcttcATGG CTGATGAAATCCAGCAT | Used for amplification of WT *OsANX9* |
| *OsANX9*-pDONR207-R | ggggaccactttgtacaagaaagctgggtcATGGC TACCAACTAGAGAAAG | Used for amplification of WT *OsANX9* |
| *OsANX10*-pDONR207-F | ggggacaagtttgtacaaaaaagcaggcttcATGG CCTCTCGGTGTCTTGT | Used for amplification of WT *OsANX10* |
| *OsANX10*-pDONR207-R | ggggaccactttgtacaagaaagctgggtcTGAGG CCCTTGACCCCTTG | Used for amplification of WT *OsANX10* |
| *mTurquoise*-qRT-F | ACCACTACCAGCAGAACACC | RT-qPCR for *mTurquoise* |
| *mTurquoise*-qRT-R | ATGTGATCGCGCTTCTCGTT | RT-qPCR for *mTurquoise* |
| *mRFP*-qRT-F | CGCCTACAAGACCGACATCA | RT-qPCR for *mRFP* |
| *mRFP*-qRT-R | CGCTCGTACTGTTCCACGAT | RT-qPCR for *mRFP* |

| Primer name | Primer sequence (5'->3') | Comments |
|---|---|---|
| *NlANX5*-nYFP-F | GACGACAAGCATTTAAATCTCGAGGGA ATGCTAGATGTTTCAGTATGTGTATT | Used for BiFC |
| *NlANX5*-nYFP-R | CATGGTGGCGATGGATCTTCTAGAGGA CTGCTGTACTCCACTGACGATAT | Used for BiFC |
| *OsANN2*-cYFP-F | GAGGACCTGCTTTCTAGACTCGAGGG AATGGCAACGATAGTAGTTCCTCC | Used for BiFC |
| *OsANN2*-cYFP-R | GATTTTGCACGCCGGACGGGTACCGG AGATGCCACTCCCAAGTAGAGC | Used for BiFC |
| *OsANN6*-cYFP-F | GAGGACCTGCTTTCTAGACTCGAGGG AATGTCCATCAATGCCGTCC | Used for BiFC |
| *OsANN6*-cYFP-R | GATTTTGCACGCCGGACGGGTACCGG AGTCCTCCCTCCCGACCAAC | Used for BiFC |
| *OsANN8*-cYFP-F | GAGGACCTGCTTTCTAGACTCGAGGG AATGGCCTCGATTAGAGATTTTG | Used for BiFC |
| *OsANN8*-cYFP-R | GATTTTGCACGCCGGACGGGTACCGG AATCTCTTGTCCTCAGGCCAACT | Used for BiFC |
| ds*NlANX5*-RNAi-F | TAATACGACTCACTATAGGGAGACCAG GTCCAGTGTCGTCAT | Used for *NlANX5* RNAi |
| ds*NlANX5*-RNAi-R | TAATACGACTCACTATAGGGAGAGGCC ATCCACATCTTTCATA | Used for *NlANX5* RNAi |
| ds*NlGFP*-RNAi-F | TAATACGACTCACTATAGGGAGAATGA GTAAAGGAGAAGAACTTTTC | Used for *NlGFP* RNAi |
| ds*NlGFP*-RNAi-F | TAATACGACTCACTATAGGGAGATTTGT ATAGTTCATCCATGCCATGT | Used for *NlGFP* RNAi |

| Primer name | Primer sequence (5’->3’) | Comments |
|---|---|---|
| *NlANX5*-qRT-F | CTGGATTTGAATCTATGGTGC | RT-qPCR for *NlANX5* |
| *NlANX5*-qRT-R | AGAGAGAGGTCTTCTGCTGGTCTGTG | RT-qPCR for *NlANX5* |
| *Nl18S*-qRT-F | CGCTACTACCGATTGAA | RT-qPCR for *Nl18S* |
| *Nl18S*-qRT-R | GGAAACCTTGTTACGACTT | RT-qPCR for *Nl18S* |
| *OsLOX1*-qRT-F | GTACGCTGGGTTCACAGCTC | RT-qPCR for *OsLOX1* |
| *OsLOX1*-qRT-R | TCAGATGGATGTGCTGTTGG | RT-qPCR for *OsLOX1* |
| *OsAOC*-qRT-F | CGTACCTGACCTACGAGGAG | RT-qPCR for *OsAOC* |
| *OsAOC*-qRT-R | GCCCTTGAGGTAGAAGGTGT | RT-qPCR for *OsAOC* |
| *OsJAR2*-qRT-F | AGAAGGTTCTCCGCCACTAC | RT-qPCR for *OsJAR2* |
| *OsJAR2*-qRT-R | CGGAGCTGAAGAAGACGTTG | RT-qPCR for *OsJAR2* |
| *OsJAmyb*-qRT-F | GAGGACCAGAGTGCAAAAGC | RT-qPCR for *OsJAmyb* |
| *OsJAmyb*-qRT-R | CATGGCATCCTTGAACCTCT | RT-qPCR for *OsJAmyb* |
| *OsJAZ9*-qRT-F | CGTCTGCGATTTGAGAATTG | RT-qPCR for *OsJAZ9* |
| *OsJAZ9*-qRT-R | ATGCGACGAGAACCATCTTC | RT-qPCR for *OsJAZ9* |
| *OsJAZ11*-qRT-F | CGTGTCTGTGGAAAGTGTGG | RT-qPCR for *OsJAZ11* |
| *OsJAZ11*-qRT-R | GCTACTAATTCCCCCGGAAG | RT-qPCR for *OsJAZ11* |
| *OsGLR6.5*-qRT-F | GGGGTCCAAGAATGGAGGTG | RT-qPCR for *OsGLR6.5* |
| *Os GLR6.5*-qRT-R | GAGGGTCCGAGACAATAGCG | RT-qPCR for *OsGLR6.5* |
| *OsGLR7.1*-qRT-F | GATTTTGCACGCCGGACGGGTACCGG | RT-qPCR for *OsGLR7.1* |
| *OsGLR7.1*-qRT-R | AATCTCTTGTCCTCAGGCCAACT | RT-qPCR for *OsGLR7.1* |
| *OsGLR7.2*-qRT-F | TGCCCTACCCCGTTTCATTC | RT-qPCR for *OsGLR7.2* |
| *OsGLR7.2*-qRT-R | TTGCAGCGTCAAAGTGGTTG | RT-qPCR for *OsGLR7.2* |
| *OsACTIN*-qRT-F | TGGACAGGTTATCACCATTGGT | RT-qPCR for *OsACTIN* |
| *OsACTIN*-qRT-R | CCGCAGCTTCCATTCCTATG | RT-qPCR for *OsACTIN* |

**Table S3. Protein sequences used for AlphaFold3.**

| Protein name | Protein sequence | UniProt BLAST |
|---|---|---|
| *Nl*ANX5 | MLDVSVCVFQNATDPPVAPVPDKPENSTGSPPAPGPVSSYK<br>PHKFLISPAQPAGVPTIFAKQVDLKQQATYFNSLISQHKLDQM<br>AIDLAKLTCEQRMQVKVEYTNAYSRSIEQDIDGKTKKTFMRLL<br>YALIQPMPELMAATIDWSITQKQDFLYVSLICTTPADLLTQIANT<br>YAAKNNKKLVDAIMQNMKDVDGQDFLSDIVNAVTGFESMVQ<br>TRPNNTISEDLIQTQVNYIPQAQGNCSEKDNPDLYKFMGGAS<br>FAQIAAVVKAFQQKTSLSAASQIRQFCNGILSEAYSRIVIFAED<br>PASYWTTEVYNSMGYGKTIDRGLTVAILYRSEIDLATIRDSFKT<br>TYHNDLVDFIKSHCSETYEKVMENIVSGVQQ | A0A220XIK6_NIL<br>LU |
| *Os*ANN1 | MATLTVPAAVPPVAEDCEQLRKAFKGWGTNEKLIISILAHRDA<br>AQRRAIRRAYAEAYGEELLRALNDEIHGKFERAVIQWTLDPAE<br>RDAVLANEEARKWHPGGRALVEIACTRTPSQLFAAKQAYHE<br>RFKRSLEEDVAAHITGDYRKLLVPLVTVYRYDGPEVNTSLAH<br>SEAKILHEKIHDKAYSDDEIIRILTTRSKAQLLATFNSYNDQFG<br>HPITKDLKADPKDEFLGTLRAIIRCFTCPDRYFEKVIRLALGG<br>MGTDENSLTRIITTRAEVDLKLIKEAYQKRNSVPLERAVAKDT<br>TRDYEDILLALLGAE | LOC_Os02g5175<br>0 |
| *Os*ANN2 | MATIVVPPVTPSPAEDADALLKAFQGWGTDEQAVIGVLAHRD<br>ATQRKQIRLTYEENYNENLIQRLQSELSGDLERAMYHWVLDP<br>VERQAVMVNTATKCIHEDYAVIVEIACTNSSSELLALLLALVST<br>YRYDGDEVNDALAKSEAKILHETVTNGDTDHGELIRIVGTRS<br>RAQLNATFSWFRDERGTSITKALQHGADPTGYSHALRTALR<br>CISDANKYFVKVLRNAMHKSGTNEDSLTRVIVLHAEKDLKGIK<br>DAFQKRASVALEKAIGNDTSGDYKSFLMALLGSGI | LOC_Os01g3127<br>0 |

| Protein name | Protein sequence | UniProt BLAST |
|---|---|---|
| *Os*ANN3 | MADEIQHLTRAFSGLGGLGVDEPAMVSALAKWRRQPEKLSGFRKSFNGFFKDHGGVIEKCEEEYMLHLAAEFSRFKNLMVMWAMHPWERDARLAHHVLHQAHPAAIVVEIACTRTAEELLGARKAYQALFHHSLEEDVAYRARDKPYCGLLVGLVSAYRYEGPRVSEETARAEAKALVAAVKSAGHAAAKLVENDDVVRILTTRSKPHLVETFKHYKEIHGRHIEEDLGHEETLREAALCLATPARYFSEVVAAAVSDGADHHAKEALTRVAVTRADVDMDAIRAAYHEQFGGRLEDAVAGKAHGYYRDALLSLVAGGK | LOC_Os05g31750 |
| *Os*ANN4 | MCCWCCCLDCIHNIPPLNLLFLHFSPHSLSSSAASAGGGEAAAAAAVAPMASISVPNPAPSPTEDAESIRKAVQGWGTDENALIEILGHRTAAQRAEIAVAYEGLYDETLLDRLHSELSGDFRSALMLWTMDPAARDAKLANEALKKKKKGELRHIWVLVEVACASSPDHLVAVRKAYRAAYASSLEEDVASCSLFGDPLRRFLVRLVSSYRYGGGGVDGELAIAEAAELHDAVVGRGQALHGDDVVRIVGTRSKAQLAVTLERYRQEHGKGIDEVLDGRRGDQLAAVLKAALWCLTSPEKHFAEVIRTSILGLGTDEEMLTRGIVSRAEVDMEKVKEEYKVRYNTTVTADVRGDTSGYYMNTLLTLVGPEK | LOC_Os05g31760 |
| *Os*ANN5 | MATLTVPSAVPPVADDCDQLRKAFQGWGTNEALIISILAHRDAAQRRAIRRAYADTYGEELLRSITDEISGDFERAVILWTLDPAERDAVLANEVARKWYPGSGSRVLVEIACARGPAQLFAVRQAYHERFKRSLEEDVAAHATGDFRKLLVPLISAYRYEGPEVNTKLAHSEAKILHEKIQHKAYGDDEIIRILTTRSKAQLIATFNRYNDEYGHPINKDLKADPKDEFLSTLRAIIRCFCCPDRYFEKVIRLAIAGMGTDENSLTRIITTRAEVDLKLITEAYQKRNSVPLERAVAGDTSGDYERMLLALLGQEQ | LOC_Os06g11800 |

| Protein name | Protein sequence | UniProt BLAST |
|---|---|---|
| *Os*ANN6 | MSINAVPSPVPSASDDAESLRKALQVRHGRMVTTRVASAGWRADKGALTRILCRRTAAQRAAIRRAYAFLYREPLLNCFRYKLSRHCLLSLDFWKAMILWTMDPAERDANLVHEALKKKQRDETYYMSVLIEMLVRLVSSYRYEGDECVVDMDVVRMEASQLAEAIKKKKQPRGEDEVVRIVTTRSKSQLRATFQRYREDHGSDIAEDIDSHCIGQFGRMLKTAVWCLTSPEKHFAEVIRHSILGLGTYEDMLTRVIVSRAEIDMRHIREEYKVRYKTTVTRDVVGDTSFGYKGFLLALVGRED | LOC_Os07g46550 |
| *Os*ANN7 | MASLSVPPVPTDPRRDAIDLHRAFKGFGCDATAVTAILAHRDASQRALIRRHYAAVYHQDLLHRLAAELSGHHKRAVLLWVLDPASRDAAVLHQALNGDVTDMRAATEVVCSRTPSQLLVVRQAYLARFGGGGGGGLEHDVAVRASGDHQRLLLAYLRSPRYEGPEVVDMAAAARDARELYRAGERRLGTDERTFIRVFSERSAAHMAAVAAAYHHMYDRSLEKAVKSETSGNFGFGLLTILRCAESPAKYFAKVLHEAMKGLGTNDTTLIRVVTTRAEVDMQYIKAEYHRSYKRSLADAVHSETSGNYRTFLLSLIGRDR | LOC_Os08g32970 |
| *Os*ANN8 | MASIRDFAKRYEADCRHLNQFFSGNVSPNNARPVLEIFTARSSQEMKQICRAYSSMYRQDLIQLLSQQKTTFAVIPASDPLVSKHISILSGSIAIRVACLRASEPCVRDADIARDALFGRRIDGDVLVEVVCTRPSGEVALIRQAYQARYSASLERDVSSRTSGSLNEVLLAFLGSSGSGYHGGRVDATMAMCDAKTLYEAVEISAARVDQRSVLQLLRHRSGDQLRAVLASYRRLYGQELARALKRKDGDTSGGGGGRRGESSSFPGILRAALRCAQLPERHFARAVRAALERGEGGAGADRRDARGRRRAPRQPGVRGQDRVDAGERRPERVRQRRHREVGRRLDRRLARGVAQVGLRTRD | LOC_Os09g20330 |

| Protein name | Protein sequence | UniProt BLAST |
|---|---|---|
| *Os*ANN9 | MASLTLPPAPTNPRQDAIDLHKAFKGFGCDSTTVINILTHRDS MQRALIQQEYRTMYSEDLSRRISSELSGHHKKAMLLWILDPA GRDATVLREALSGDTIDLRAATEIICSRTPSQLQIMKQTYHAK FGTYLEHDIGQRTSGDHQKLLLAYVGIPRYEGPEVDPTIVTHD AKDLYKAGEKRLGTDEKTFIRIFTERSWAHMASVASAYHHMY DRSLEKVVKSETSGNFELALLTILRCAENPAKYFAKVLRKSMK GMGTDDSTLIRVVVTRTEIDMQYIKAEYYKKYKKSLAEAIHSE TSGNYRTFLLSLVGSH | LOC_Os09g2316 0 |
| *Os*ANN10 | MASRCLVTTGFEDECREIHDACNQPRRLSVLLAHRSPSERQ KIKATYRTVFGEDLAGEVQKILMVNQEDELCKLLYLWVLDPSE RDAIMARDAVENGGATDYRVLVEIFTRRKQNQLFFTNQAYLA RFKKNLEQDMVTEPSHPYQRLLVALATSHKSHHDELSRHIAK CDARRLYDAKNSGMGSVDEAVILEMFSKRSIPQLRLAFCSYK HIYGHDYTKALKKNGFGEFEQSLRVVVKCIYNPSMYFSKLLH RSLQCSATNKRLVTRAILGSDDVDMDKIKSVFKSSYGKDLED FILESLPENDYRDFLLGAAKGSRAS | LOC_Os09g2799 0 |
